# A sex-specific switch in a single glial cell patterns the apical extracellular matrix

**DOI:** 10.1101/2023.03.17.533199

**Authors:** Wendy Fung, Taralyn M. Tan, Irina Kolotuev, Maxwell G. Heiman

## Abstract

Apical extracellular matrix (aECM) constitutes the interface between every tissue and the outside world. It is patterned into diverse tissue-specific structures through unknown mechanisms. Here, we show that a male-specific genetic switch in a single *C. elegans* glial cell patterns the aECM into a ∼200 nm pore, allowing a male sensory neuron to access the environment. We find that this glial sex difference is controlled by factors shared with neurons (*mab-3, lep-2, lep-5*) as well as previously unidentified regulators whose effects may be glia-specific (*nfya-1, bed-3, jmjd-3.1*). The switch results in male-specific expression of a Hedgehog-related protein, GRL-18, that we discover localizes to transient nanoscale rings at sites of aECM pore formation. Blocking male-specific gene expression in glia prevents pore formation, whereas forcing male-specific expression induces an ectopic pore. Thus, a switch in gene expression in a single cell is necessary and sufficient to pattern aECM into a specific structure.

## INTRODUCTION

The apical extracellular matrix (aECM) is a conserved, intricate network of secreted macromolecules that lines the outward or luminal-facing surfaces of epithelial tissues, including the lungs, vasculature, and gut, as well as sense organs like the olfactory epithelium and inner ear (Ando et al., 2019; Gaudette et al., 2020; Goodyear and Richardson, 2018; Johansson et al., 2013; Whitsett et al., 2015). The aECM is often viewed as a static protective barrier, but growing evidence indicates that it is dynamic during development, diverse in composition and form, and plays crucial roles in tissue morphogenesis (Li Zheng et al., 2020). In addition to providing a defense against pathogens and desiccation, distinct aECM structures establish and maintain epithelial tubes, control the function of sense organs by forming nanopores for odor reception or resonating membranes for hearing, and shape organs through the distribution of tensile forces (Ando et al., 2019; Dong and Hayashi, 2015; Gaudette et al., 2020; Gill et al., 2016; Goodyear and Richardson, 2018; Johansson et al., 2013; Whitsett et al., 2015). The structure of aECM is thus tailored to the needs of a particular tissue or organ. However, the mechanisms that pattern aECM remain largely mysterious.

The *C. elegans* cuticle is an aECM layer that coats the surface of the animal and provides a model of aECM patterning. As animals transition through four juvenile (larval) stages to the adult stage, they undergo molts in which they shed the old cuticle and form a new cuticle that is specialized for each developmental stage (Page, 2007). We have focused on intricate aECM specializations that are associated with sense organs in the head, midbody, and tail (Ward et al., 1975). Each sense organ contains one or more sensory neurons with long unbranched dendrites that terminate in sensory cilia, as well as two glial cells called the sheath and socket. The sheath glial cell wraps the dendrite endings, while the socket glial cell forms a cellular pore in the skin through which neuronal sensory cilia protrude. This cellular pore is overlaid by cuticle aECM that takes the form of either a closed sheet or an open pore, depending on the function of the sense organ (Figure 1A). A closed sheet of aECM is found overlying sense organs that contain a single mechanosensory neuron, whose cilium is embedded directly in the cuticle to sense external forces (Figure 1A, left). By comparison, open pores in the cuticle are found at bifunctional sense organs that contain both a mechanosensory and a chemosensory neuron (Figure 1A, right). In these sense organs, the chemosensory cilium protrudes through the open pore in the cuticle in order to directly access chemical cues in the external environment.

**Figure 1.**
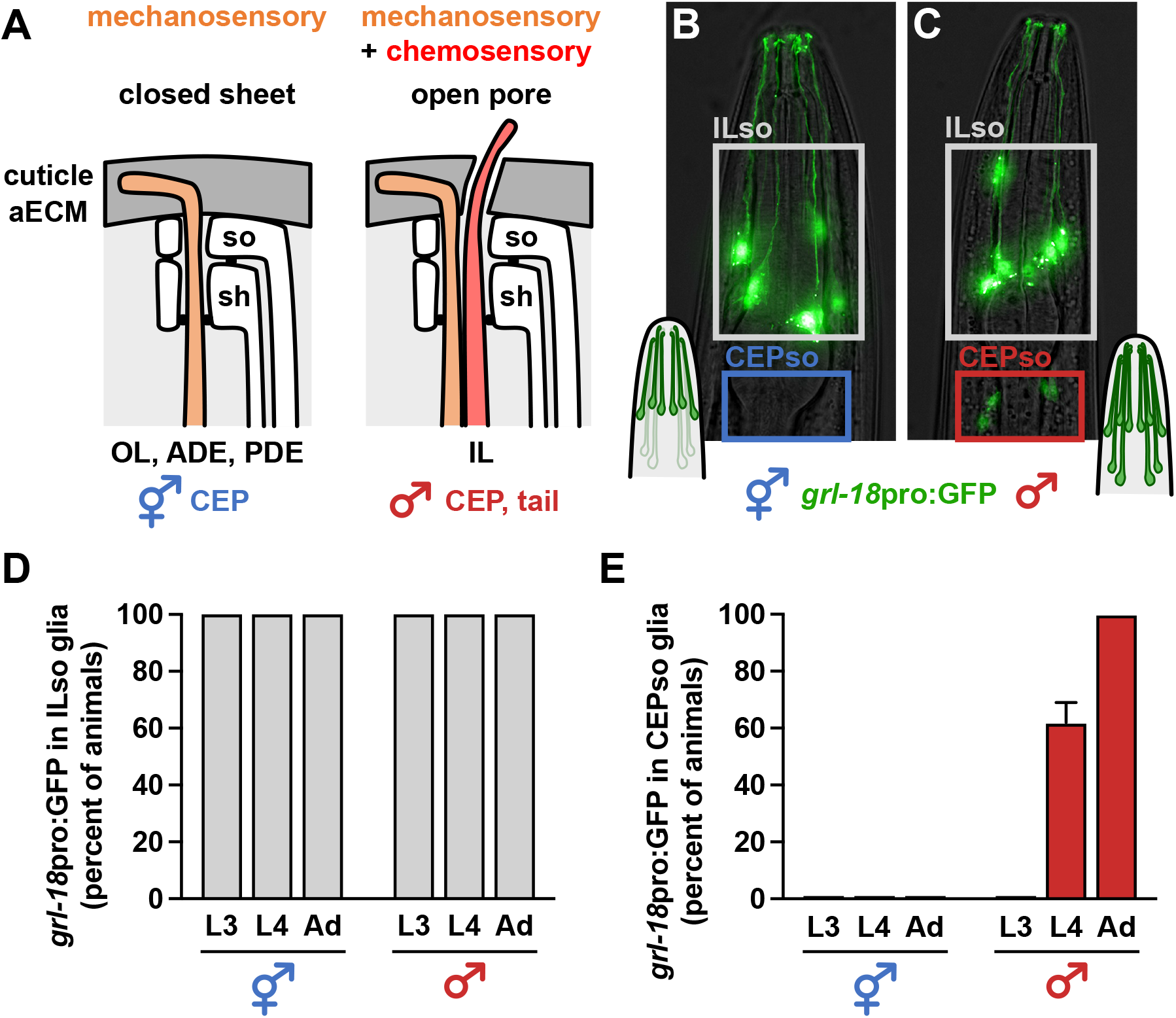
Glia initiate male-specific gene expression at sexual maturity. (**A**) Schematic of *C. elegans* sense organs showing socket (so) and sheath (sh) glial cells wrapping neuronal endings. Cuticle aECM forms a closed sheet (left) over sense organs with a single mechanosensory neuron (outer labial, OL; deirids, ADE/PDE) or a ∼200 nm open pore (right) over sense organs with a mechanosensory and chemosensory neuron pair (inner labial, IL; rays and hook of male tail). Notably, cuticle aECM of the cephalic (CEP) sense organ undergoes sex-specific remodeling from a closed sheet in hermaphrodites to an open pore in adult males. (**B, C**) Expression of *grl-18*pro:GFP in (B) hermaphrodite and (C) male head glia. Nose is up. (**D, E**) Fraction of hermaphrodites (n=48) and males (n=50) expressing *grl-18*pro:GFP at third and fourth larval stages (L3, L4) and the 1-day adult stage (Ad) in (D) ILso glia and (E) CEPso glia. The same animals were followed and scored at each stage. Error bars, standard error of the mean (SEM).

To better understand how specialized aECM is patterned, we focused on a discrete sex-specific aECM remodeling event associated with the four cephalic (CEP) sense organs in the head. In hermaphrodites, each CEP sense organ contains only a single mechanosensory neuron called CEP and the overlying cuticle forms a closed sheet (Figure 1A; orange, CEP). By contrast, in adult males, each CEP sense organ contains an additional chemosensory neuron called CEM and the overlying cuticle forms a ∼200 nm open pore through which the CEM cilium protrudes to detect pheromones from mating partners (Chasnov et al., 2007; Srinivasan et al., 2008; Ward et al., 1975; White et al., 2007) and to release extracellular vesicles thought to mediate social communication (Wang et al., 2014) (Figure 1A; red, CEM). The CEM neurons undergo embryonic apoptosis in hermaphrodites (Sulston et al., 1983) but, in males, they differentiate and form a mature sensory cilium at the onset of sexual maturation in the fourth larval (L4) stage (Akella et al., 2019). Thus, the cuticle remodels from a closed sheet to an open pore during the L4/adult molt in males. It was previously inferred that *“the openings in males must be created by the CEM dendrites*,*”* (Perkins et al., 1986) because they appeared to be the only cells that differ between the sexes in this sense organ. Surprisingly, we find that one of the glial cells also differs between the sexes, switching on a gene expression program at the L4/adult molt only in males. Remarkably, this glial switch is necessary and sufficient to create the cuticle pore. Overall, we identify a novel sexual dimorphism in a single glial cell, define its upstream regulators, and show that a switch in gene expression in a single cell is enough to induce dramatic remodeling of the aECM.

## RESULTS

### Glia undergo a sex-specific switch in gene expression

We serendipitously found that CEP socket (CEPso) glia undergo a sex-specific switch in gene expression. Previously, we showed that a 2968-bp transcriptional reporter for the putative secreted protein GRL-18 (*grl-18*pro:GFP, Figure S1A) is a cell-type specific marker for IL socket (ILso) glia in hermaphrodites (Figure 1B) (Cebul et al., 2020; Fung et al., 2020). Using this transgene or a reporter engineered at the endogenous locus (*grl-18*-SL2-YFP:H2B, Figure S1F), we observed expression in ILso glia in both sexes throughout life, as well as in sex-specific reproductive structures – the hermaphrodite vulval epithelial cells and male tail glia – shortly after these cells are born at the L4 stage (Figures 1B-1D and Figures S1B-S1J). Strikingly, we also observed expression in CEPso glia in L4 and adult males, but never in hermaphrodites, indicating that these glia are sexually dimorphic (Figures 1B-1C, 1E and Figures S1G, S1I). Thus, CEPso glia undergo a sex-specific switch in gene expression that coincides with remodeling of the cuticle aECM from a sheet to a pore.

While sexual dimorphism is well-established in *C. elegans* neurons, only two examples have been reported in glia, both involving the production of male-specific neurons rather than altered function of the glial cell itself (Molina-García et al., 2020; Sammut et al., 2015). CEPso glia develop embryonically, but our reporter suggests that a sexually dimorphic switch in gene expression occurs specifically at the L4 stage. Due to possible perdurance of the fluorescent reporter, we cannot infer whether this switch is transient or persists throughout adulthood. Notably, there is a nearly perfect correspondence between the glia that express this reporter and those that are associated with narrow cuticle pores (ILso, CEPso, male tail glia) (Perkins et al., 1986; Sulston et al., 1980; Ward et al., 1975), suggesting a shared gene expression program in these glial types.

### The sex-specific switch in gene expression is controlled cell autonomously in glia

Sexually dimorphic CEPso glial gene expression could be induced cell autonomously or non-cell autonomously, for example via male-specific signals from CEM or other neurons. To distinguish these possibilities, we genetically altered the sex identity of neurons or glia through cell-type-specific mis-expression of the masculinizing factor *fem-3* or the feminizing factor *tra-2*(IC) from the sex determination pathway of *C. elegans*, as previously described (Lee and Portman, 2007; Mehra et al., 1999; Sammut et al., 2015; White et al., 2007) (Figures 2A, 2E). Neurons and glia were targeted using a pan-neuronal promoter (*rab-3*pro) (White et al., 2007), a pan-glial promoter (*mir-228*pro) (Pierce et al., 2008), and a newly identified CEPso-specific promoter (*col-56*pro, Figure S2). We found that masculinization of glia induced inappropriate *grl-18* expression in hermaphrodite CEPso glia, whereas feminization of glia blocked *grl-18* expression in male CEPso glia (Figures 2B, 2F). In contrast, sex-reversal of neurons had no effect (Figures 2B, 2F).

**Figure 2.**
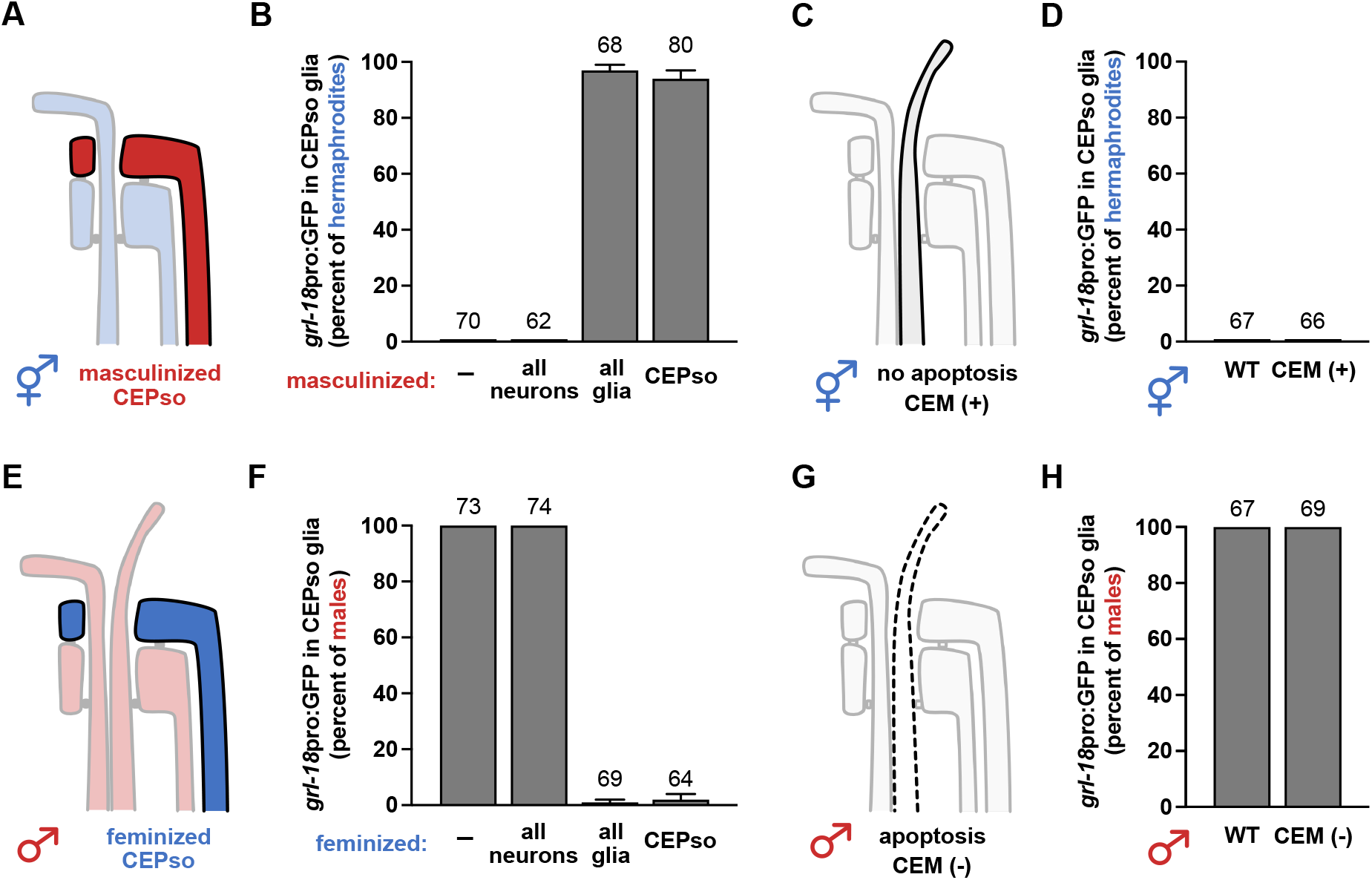
The sex-specific switch in gene expression is controlled cell autonomously in glia and does not depend on male neurons. **(A-B, E-F)** Neurons or glia were (A) masculinized in hermaphrodites or (E) feminized in males via mis-expression of *fem-3* or *tra-2*(IC), respectively, using cell-type-specific promoters (all neurons, *rab-3*; all glia, *mir-228*; CEPso glial, *col-56*, see Figure S2). Fraction of 1-day adult (B) hermaphrodites or (F) males that express *grl-18*pro:GFP in CEPso glia is shown. **(C-D, G-H)** To test if CEM neurons are sufficient and/or necessary for the switch, *grl-18*pro:GFP expression was evaluated in (C) *ceh-30(3714gf)* hermaphrodites in which CEM neurons inappropriately survive (CEM+), and (G) in *ceh-30(4289lf)* males in which CEM neurons inappropriately undergo apoptosis (CEM-) (Schwartz and Horvitz, 2007). Fraction of 1-day adult (D) CEM+ hermaphrodites and (H) CEM-males that express *grl-18*pro:GFP in CEPso glia. Sample sizes are indicated above the bars of each graph. Error bars, SEM.

We also directly tested whether CEM neurons contribute to the change in glial gene expression by taking advantage of *ceh-30* loss-of-function (*lf*) or gain-of-function (*gf*) mutants in which CEM neurons inappropriately die in males or survive in hermaphrodites, respectively. *ceh-30* encodes a Bar family homeodomain transcription factor that is normally expressed in male CEM neurons and promotes their survival (Peden et al., 2007; Schwartz and Horvitz, 2007). In the *ceh-30(lf)* mutant, most of the homeodomain sequence is deleted and the transcription factor is non-functional; whereas in the *ceh-30(gf)* mutant, a *cis*-regulatory binding site for the sex determination factor TRA-1 is disrupted, leading to mis-expression of active CEH-30 protein in hermaphrodites (Schwartz and Horvitz, 2007) (Figure 2C, 2G). We observed that *grl-18* expression in CEPso glia remains unaffected in *ceh-30(lf)* males and *ceh-30(gf)* hermaphrodites (Figures 2D, 2H), showing that CEM neurons are neither necessary nor sufficient for the switch in glial gene expression.

Together, these results suggest that sexually dimorphic gene expression in the glia is controlled cell autonomously by the sex identity of the glial cell itself.

### Regulators of sex- and timing-dependent changes in glial gene expression

To identify the regulatory factors that control sexual dimorphism in glia, we used forward and candidate-based screens to isolate mutants that exhibit delayed or absent *grl-18* expression in males (OFF mutants; Figure 3A, left), or inappropriate *grl-18* expression in hermaphrodites (ON mutants; Figure 3A, right). We identified three classes of genes that regulate the sex-specificity and timing of glial gene expression (Table 1 and Table S1): DM domain transcription factors (class I), heterochronic genes (class II), and novel regulators (class III).

**Figure 3.**
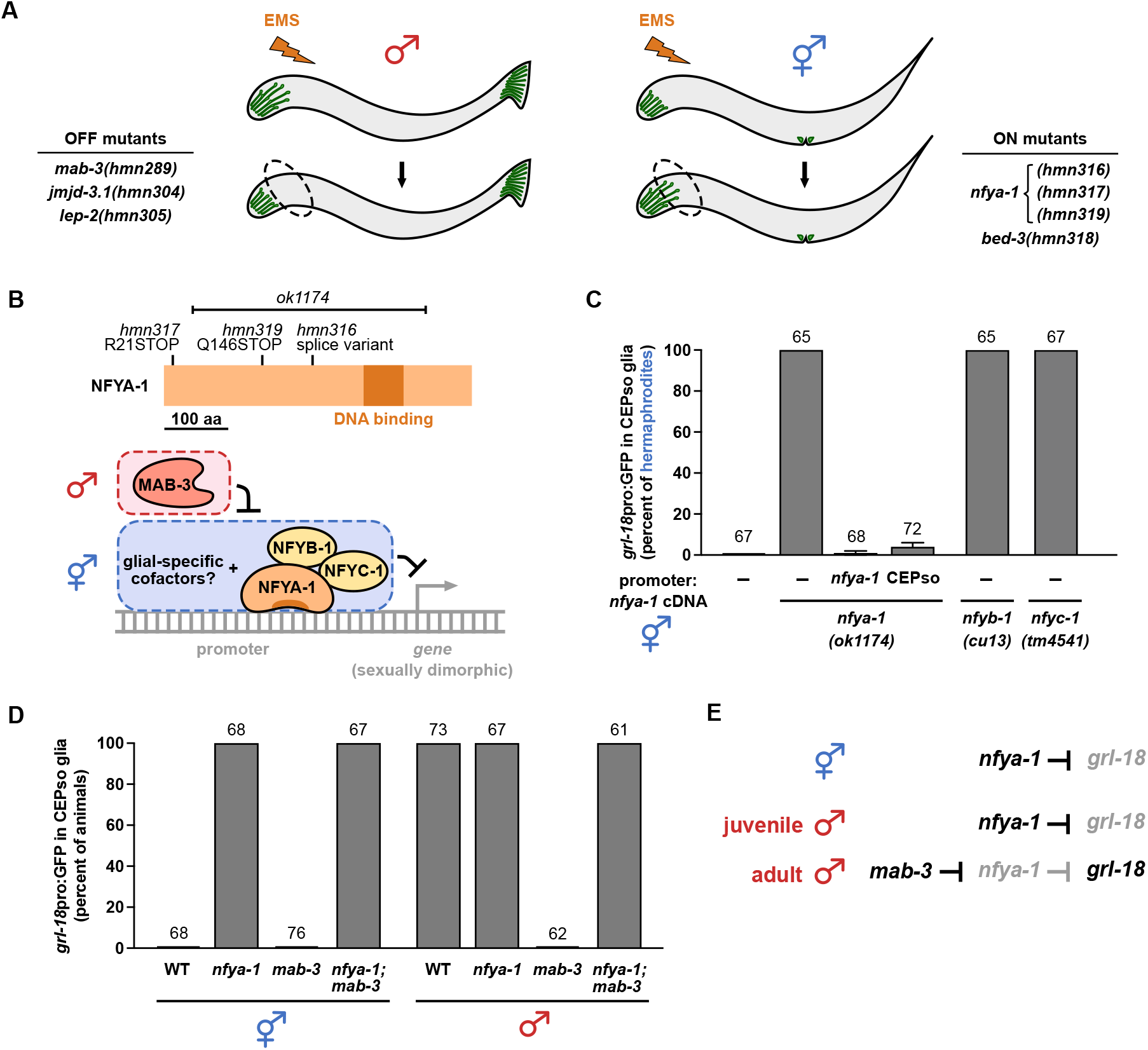
Sex-specific glial gene expression requires the NF-Y repressive complex acting in the glial cell downstream of MAB-3. **(A)** Schematic of forward genetic screens in which animals expressing *grl-18* reporters were mutagenized with ethyl methanesulfonate (EMS). Mutant males lacking *grl-18* expression in CEPso glia (OFF mutants, left) and mutant hermaphrodites exhibiting inappropriate *grl-18* expression in CEPso glia (ON mutants, right) were isolated, and the causal mutations were identified (see Table 1). **(B)** Schematic of NFYA-1 protein showing effects of novel alleles (*hmn316*, *hmn317*, *hmn319*) and an existing 1837 bp deletion (*ok1174*). Working model of the NF-Y repressor complex (NFYA-1, NFYB-1, NFYC-1) acting with glial-specific cofactors to inhibit male-specific gene expression, subject to de-repression by MAB-3. (**C**) Fraction of 1-day adult wild-type or *nfya-1*, *nfyb-1*, or *nfyc-1* mutant hermaphrodites that express *grl-18*pro:GFP in CEPso glia. *nfya-1* mutant is shown alone (–) or with rescuing *nfya-1* cDNA under control of *nfya-1* or CEPso-specific (*col-56*) promoters. (**D**) Epistasis analysis showing fraction of 1-day adult *mab-3(mu15)*, *nfya-1(ok1174)*, or *mab-3(mu15); nfya-1(ok1174)* mutants of each sex that express *grl-18*pro:GFP in CEPso glia. (**E**) Genetic model of *grl-18* regulation in CEPso glia. Sample sizes are above the bars of each graph. Error bars, SEM.

**Table 1.**
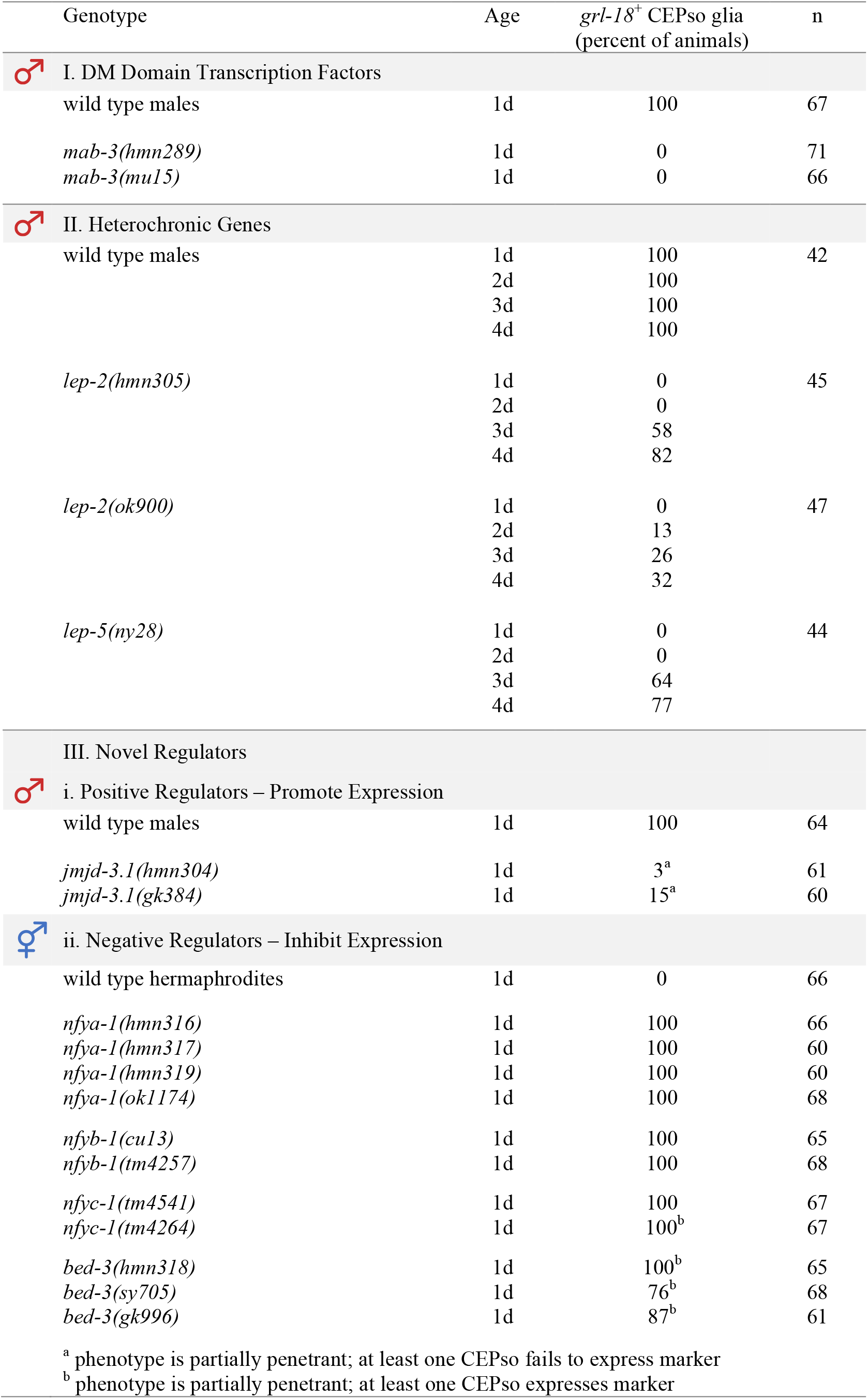
Regulators of sex-specific glial gene expression. Percent of males (I, II, III(i)) or hermaphrodites (III(ii)) at the indicated adult age that express *grl-18*pro:mApple or *grl-18*pro:GFP in CEPso glia. Expression in ILso glia and male tail glia were unaffected in all cases. Based on the sequence changes involved, the *hmn304* and *gk384* alleles of *jmjd-3.1*, the *tm4264* allele of *nfyc-1*, and the *hmn318*, *sy705*, and *gk996* alleles of *bed-3* are unlikely to be nulls.

| Genotype | Age | *grl-18<sup>+</sup>* CEPso glia<br>(percent of animals) | n |
| --- | --- | --- | --- |
| ♂ I. DM Domain Transcription Factors |  |  |  |
| wild type males | 1d | 100 | 67 |
| *mab-3(hmn289)* | 1d | 0 | 71 |
| *mab-3(mu15)* | 1d | 0 | 66 |
| ♂ II. Heterochronic Genes |  |  |  |
| wild type males | 1d | 100 | 42 |
|  | 2d | 100 |  |
|  | 3d | 100 |  |
|  | 4d | 100 |  |
| *lep-2(hmn305)* | 1d | 0 | 45 |
|  | 2d | 0 |  |
|  | 3d | 58 |  |
|  | 4d | 82 |  |
| *lep-2(ok900)* | 1d | 0 | 47 |
|  | 2d | 13 |  |
|  | 3d | 26 |  |
|  | 4d | 32 |  |
| *lep-5(ny28)* | 1d | 0 | 44 |
|  | 2d | 0 |  |
|  | 3d | 64 |  |
|  | 4d | 77 |  |
| III. Novel Regulators |  |  |  |
| ♂ i. Positive Regulators – Promote Expression |  |  |  |
| wild type males | 1d | 100 | 64 |
| *jmjd-3.1(hmn304)* | 1d | 3<sup>a</sup> | 61 |
| *jmjd-3.1(gk384)* | 1d | 15<sup>a</sup> | 60 |
| ♀ ii. Negative Regulators – Inhibit Expression |  |  |  |
| wild type hermaphrodites | 1d | 0 | 66 |
| *nfya-1(hmn316)* | 1d | 100 | 66 |
| *nfya-1(hmn317)* | 1d | 100 | 60 |
| *nfya-1(hmn319)* | 1d | 100 | 60 |
| *nfya-1(ok1174)* | 1d | 100 | 68 |
| *nfyb-1(cu13)* | 1d | 100 | 65 |
| *nfyb-1(tm4257)* | 1d | 100 | 68 |
| *nfyc-1(tm4541)* | 1d | 100 | 67 |
| *nfyc-1(tm4264)* | 1d | 100<sup>b</sup> | 67 |
| *bed-3(hmn318)* | 1d | 100<sup>b</sup> | 65 |
| *bed-3(sy705)* | 1d | 76<sup>b</sup> | 68 |
| *bed-3(gk996)* | 1d | 87<sup>b</sup> | 61 |

<sup>a</sup> phenotype is partially penetrant; at least one CEPso fails to express marker<sup>b</sup> phenotype is partially penetrant; at least one CEPso expresses marker

Class I – We determined that sex-specificity is controlled by the DM domain transcription factor *mab-3*/DMRT, a conserved protein known to establish sexual dimorphism in vertebrates and invertebrates (Burtis and Baker, 1989; Raymond et al., 1998; Raymond et al., 2000). In *C. elegans*, *mab-3* is required for promoting male sexual differentiation in neurons and hypodermal cells of the tail (Fagan et al., 2018; Mason et al., 2008; Ross et al., 2005; Shen and Hodgkin, 1988; Yi et al., 2000). From a screen for mutant males lacking *grl-18* expression in CEPso glia (Figure 3A, left), we isolated a R90C mutation in the second DNA-binding domain of MAB-3 (*hmn289*) and confirmed this phenotype using a previously characterized *mab-3* null allele (*mu15*) (Raymond et al., 1998). This phenotype is fully penetrant (Table 1). Mutations in all other DM domain transcription factors of *C. elegans* were also assessed (except *dmd-4*, which is lethal) and found to have no effect (Table S1). This result shows that sex identity in glia is controlled by the same upstream regulator that is used in neurons (Fagan et al., 2018; Ross et al., 2005; Yi et al., 2000).

Class II – Heterochronic genes are conserved regulatory factors that control developmental timing and sexual development (Abreu et al., 2015; Lawson et al., 2019; Moss, 2007; Pereira et al., 2019). We found that male-specific *grl-18* expression is absent or delayed in mutants lacking the timing factors *lep-2*/Makorin (International *C. elegans* Gene Knockout Consortium) and *lep-5*, a long non-coding RNA (Kiontke et al., 2019) (Table 1). In the screen described above (Figure 3A, left), we isolated a new allele of *lep-2(hmn305)* that exhibits the same delayed expression phenotype (Table 1). This result shows that the onset of sexual maturation in glia is triggered by the same timing factors that are used in neurons (Lawson et al., 2019).

Class III – Finally, we identified novel regulators of sex-specific gene expression in glia that had not been identified in studies of neuronal sex differences. Mutations that disrupt the histone-modifier *jmjd-3.1* and the transcription factor *bed-3* result in incompletely penetrant defects, where *jmjd-3.1* mutants fail to initiate *grl-18* expression in male CEPso glia and *bed-3* mutants inappropriately initiate *grl-18* expression in hermaphrodite CEPso glia (Table 1). Most notably, we isolated three alleles of *nfya-1* (*hmn316*, *hmn317*, *hmn319*) that result in inappropriate male-like expression of *grl-18* in hermaphrodite CEPso glia (Figure 3A, right and Table 1). The phenotype is completely penetrant in all three alleles, as well as in an existing allele (*ok1174*) (International *C. elegans* Gene Knockout Consortium, Table 1). Together, these results suggest that sex identity in glia may be controlled by some factors that are distinct from those used by neurons.

### NFYA-1 is a repressor that acts downstream of MAB-3

We chose to focus on *nfya-1* because it showed the strongest phenotype. In *C. elegans*, loss of *nfya-1* has previously been shown to affect vulva development, sensory neuron specification, and expression of specific cell fate transcription factors (Deng et al., 2007; Milton et al., 2013; Aklilu et al., 2022; Heo et al., 2022). NFYA-1/NFY-A is a subunit of the conserved, ubiquitously expressed trimeric repressor complex NF-Y/CBF. It associates with the NFYB-1/NFYC-1 dimer and contains the DNA-binding domain of the regulatory complex (Deng et al., 2007; Dolfini et al., 2012). All four fully penetrant alleles of *nfya-1* carry mutations that are predicted to generate a truncated protein lacking the DNA-binding domain (Figure 3B). The *nfya-1* phenotype is rescued when *nfya-1* cDNA is expressed using the CEPso-specific promoter, demonstrating that *nfya-1* acts cell autonomously to prevent male gene expression in CEPso glia (Figure 3C). *nfyb-1* and *nfyc-1* mutants exhibit the same completely penetrant phenotype as *nfya-1* mutants, suggesting the entire complex is required for regulating sex-specific glial gene expression (Figure 3C and Table 1).

We considered the possibility that inappropriate expression of *grl-18* in these mutants simply reflects a general defect in transcriptional repression. However, two lines of evidence argue against this. First, we found that loss of *nfya-1* does not affect known sex differences in neurons (Ryan et al., 2014; Wexler et al., 2020) or other glia (Sammut et al., 2015) (Figure S3). Second, while *nfya-1*, *nfyb-1*, or *nfyc-1* mutant hermaphrodites inappropriately express *grl-18* in CEPso glia, they do so only at sexual maturity in the L4/adult stages, suggesting that repression of this gene in juveniles remains intact. Together, these results suggest that the phenotype we observe is specific to loss of sex identity of the CEPso glia.

Interestingly, although both MAB-3 and NFYA-1 are transcriptional repressors, they act in opposite directions to regulate glial gene expression: *mab-3* promotes the switch to male-specific expression, whereas *nfya-1* prevents this switch. To place them in a genetic pathway, we performed epistasis analysis. We observed that *mab-3*; *nfya-1* males switch on *grl-18* with complete penetrance, showing that loss of *nfya-1* fully bypasses the requirement for *mab-3* (Figure 3D). This leads to the model that, in hermaphrodites and juvenile males, NFYA-1 prevents *grl-18* expression in CEPso glia, whereas in sexually mature males, MAB-3 switches off NFYA-1-dependent repression, thus allowing *grl-18* to be expressed (Figure 3B, 3E). This model therefore places NFYA-1 genetically downstream of MAB-3.

It is important to note that MAB-3 may not directly act on the NFYA-1 repressive complex (which is expressed broadly in many cell types (Deng et al., 2007)) but rather on glial-specific cofactors that may recruit NFYA-1 to cell-type-specific promoters (diagrammed in Figure 3B). Similarly, the NFYA-1 complex may not act directly on *grl-18* but instead may control expression of an unidentified activator that then switches on a battery of male-specific glial genes, including *grl-18*. While it will be important to determine these mechanistic details, MAB-3 and NFYA-1 provide entry points to defining the genetic control of sexual dimorphism in glia.

### GRL-18 is a novel aECM component that localizes to transient nanoscale rings during cuticle patterning

To assess possible functions of the male-specific switch in CEPso glial gene expression, we examined localization of the GRL-18 protein. Using CRISPR/Cas9 genome editing, a sfGFP tag was inserted after the GRL-18 signal sequence at its endogenous locus (Table S4). We observed that sfGFP-GRL-18 protein appears to localize in the cuticle, forming distinctive rings at the nose tip, corresponding to the positions of the IL and male CEP sense organs (Figure 4A). Similar rings were seen at sense organ endings in the developing male tail (Figure 4B). Notably, each of these head and tail sense organs contains a single chemosensory neuron that protrudes through a narrow cuticle pore in the adult animal, suggesting that GRL-18 marks the sites of cuticle pores.

**Figure 4.**
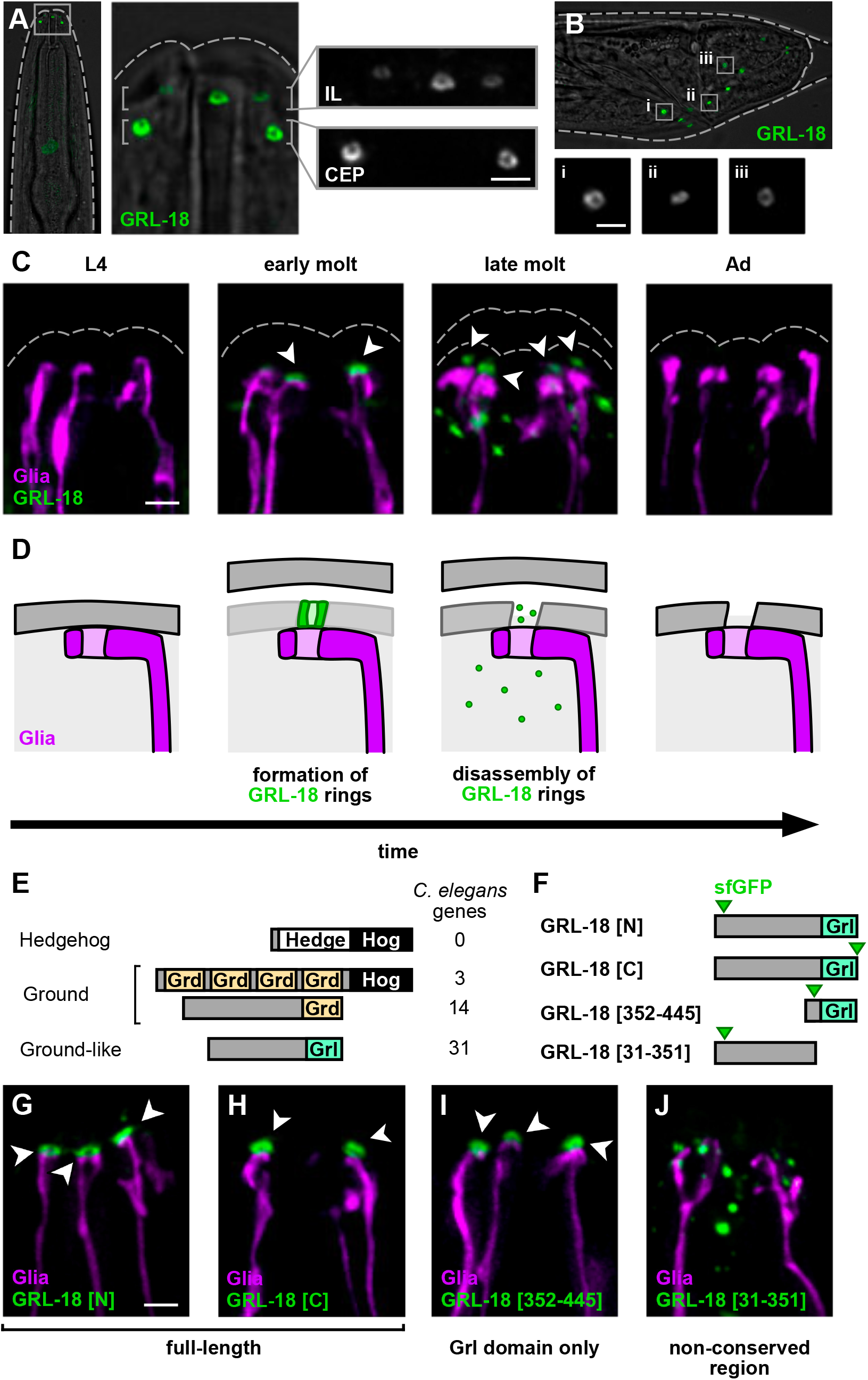
GRL-18 is a novel aECM component that transiently localizes to nanoscale rings. **(A)** Endogenously-tagged full-length sfGFP-GRL-18 forms rings at the endings of the IL and CEP sense organs in the male head. Views are maximum intensity projections through half of the head nearest the objective, showing three of the six IL sense organs and two of the four CEP sense organs; dashed lines, outline of cuticle. **(B)** Endogenously-tagged full-length sfGFP-GRL-18 rings are also found at the ray sense organs of the developing male tail during the L4/adult mid-molt stage; dashed lines, outline of cuticle. Insets (**i-iii**), magnification of rings from the boxed regions. **(C)** Localization of endogenously-tagged full-length sfGFP-GRL-18 in males through the L4 to adult (Ad) transition, showing early and late molt stages (see Figure S4 for quantification of each localization pattern). Green, sfGFP-GRL-18; magenta, glial endings; dashed lines, outline of cuticle. Arrowheads, sfGFP-GRL-18 rings. **(D)** Schematic model showing transient localization of GRL-18 to developing aECM pores during patterning of the new cuticle. **(E)** *C. elegans* gene families distantly related to Hedgehog include *ground* (*grd*) and *ground-like* (*grl*) genes. **(F)** Diagram of GRL-18 full-length proteins and fragments containing conserved C-terminal Grl domain [352-445] or non-conserved N-terminal region [31-351] expressed as low-copy transgenes under control of the *grl-18* promoter. sfGFP was inserted after the predicted signal sequence or at the C-terminus (green arrows). **(G-J)** Localization of sfGFP-GRL-18 at nose tips of L4/adult mid-molt males expressing (G) sfGFP-GRL-18, (H) GRL-18-sfGFP, (I) sfGFP-Grl domain [352-445], and (J) sfGFP-N-terminal region [31-351] transgenes. Green, sfGFP-GRL-18; magenta, glial endings. Arrowheads, sfGFP-GRL-18 rings. All scale bars, 2 µm.

We further determined that these sfGFP-GRL-18 rings are transient, forming adjacent to glial endings at the onset of the L4/adult molt and disassembling within ∼1-4 hours as the newly synthesized cuticle is complete and shedding of the old cuticle begins (Figure 4C and Figure S4). The transient localization of GRL-18 is reminiscent of the provisional matrix formed by other *C. elegans* aECM components during molts to pattern the newly synthesized cuticle (Katz et al., 2022). Provisional matrix components are thought to provide a temporary structural template or scaffold that is critical for patterning newly synthesized aECM, but they are not stably incorporated into the mature structure (Cohen et al., 2020; Gill et al., 2016; Katz et al., 2022; Öztürk-Çolak et al., 2016a). Together, our results suggest that GRL-18 is a novel aECM component that localizes to transient rings at sites where cuticle pores will form (Figure 4D).

Next, we considered what sequences in GRL-18 are required for this localization. *grl-18* belongs to a family of *ground-like* (*grl*) genes with distant homology to Hedgehog (Aspöck et al., 1999; Bürglin, 1996; Hao et al., 2006) (Figure 4E). Specifically, Hedgehog has an N-terminal ‘Hedge’ signaling domain and a C-terminal ‘Hog’ catalytic domain that is removed prior to secretion. *C. elegans* lacks Hedgehog proteins but contains several gene families with a Hog domain, for example Groundhog (Grd) (Figure 4E; also Warthog, Quahog, and Hog, not shown). *C. elegans* also contains genes that encode a Grd or Grd-like (Grl) domain but lack a Hog domain (Figure 4E). *grl-18* is one of 31 Grl domain-containing proteins in the genome, none of which have been assigned a function.

In contrast to the expectation that Grl proteins are diffusible Hedgehog-like signaling molecules, our finding that GRL-18 appears to be a structural aECM component led us to consider the role of the Grl domain in its localization. To investigate this, we expressed low-copy transgenes consisting of full-length GRL-18 or fragments with or without the Grl domain (Figure 4F). We found that either N- or C-terminally tagged full-length GRL-18 localizes to rings consistent with the endogenously-tagged protein (Figure 4G-4H; compare Figure 4C), suggesting GRL-18 is unlikely to undergo internal cleavage. The 93-amino acid C-terminal Grl domain localized identically to the full-length protein (Figure 4F, 4I), whereas the non-conserved N-terminal region localized in diffuse puncta and was not observed in rings (Figure 4F, 4J). This demonstrates that the conserved Grl domain is required for localization to nanoscale rings. It is intriguing to consider whether other Grl domain proteins are also components of the aECM, possibly reflecting an ancestral role for Hedgehog-related proteins in patterning aECM structures.

### Sex-specific changes in glial gene expression remodel the aECM

The male-specific expression of GRL-18 and its localization to cuticle rings led us to wonder if male-specific glial gene expression is required to pattern a cuticle pore for the CEM cilium. To test this possibility, first we assessed the trajectory of CEM chemosensory cilia in animals with genetically feminized CEPso glia. In wild-type males, CEM chemosensory cilia bend outwards and their tips poke past the cuticle, whereas CEP mechanosensory cilia bend inwards and track along the cuticle (Figures 5A-5B). However, in males with feminized glia, the CEM endings fail to project outwards normally and instead appear to bend inwards, in the same direction as the CEP endings (Figures 5A, 5C), suggesting that cuticle pores may not have properly formed (Figure 5D).

**Figure 5.**
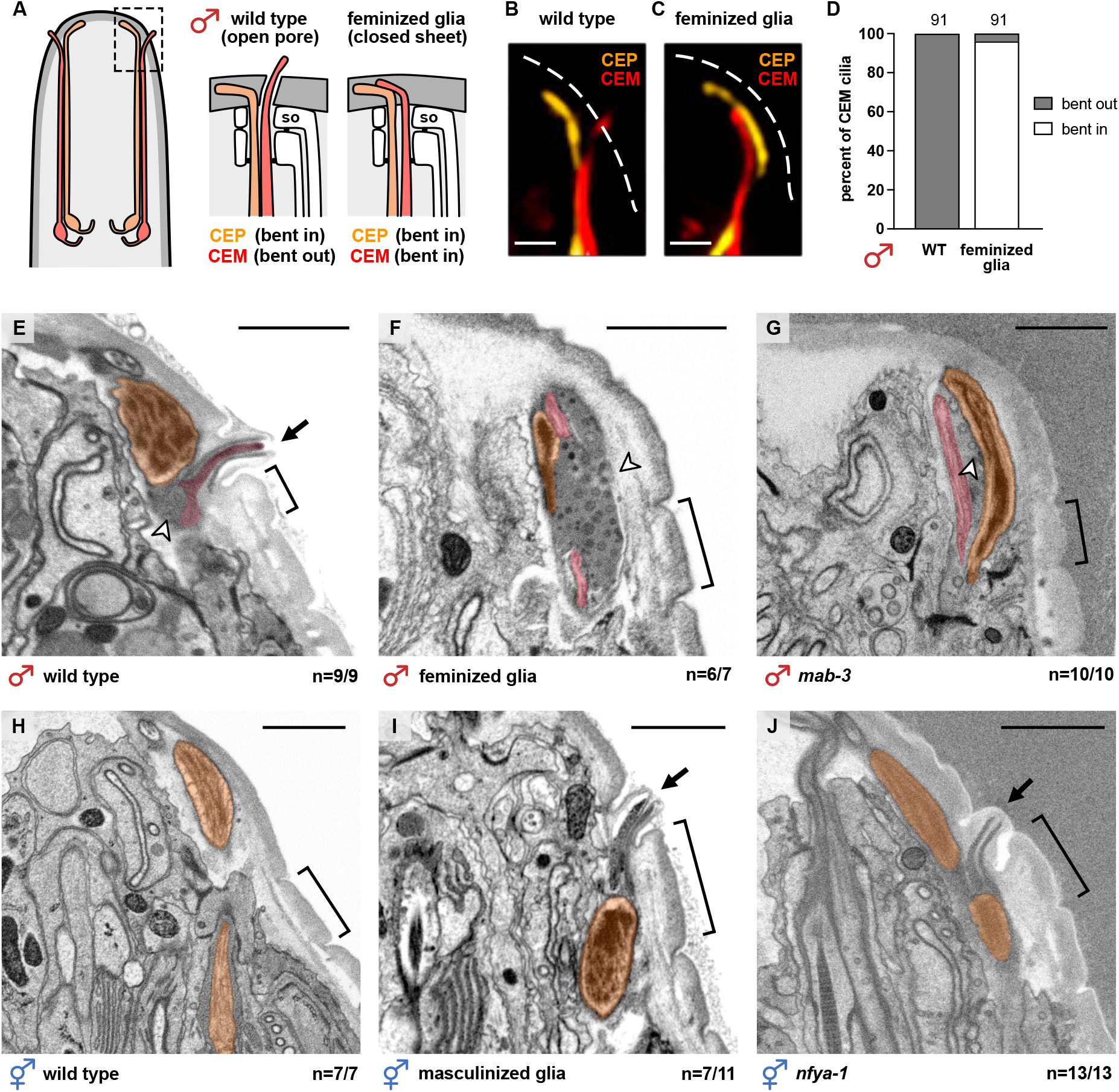
Male-specific glial gene expression is necessary and sufficient to form an aECM pore. (**A-C**) Schematic and images of CEP (orange) and CEM (red) cilia in 1-day adult males, either (B) wild-type or (C) with feminized glia (*col-56*pro:*tra-2*(IC)). Dashed white lines, outline of cuticle; scale bars, 1 µm. (**D**) Binary scoring of CEM cilia trajectories. (**E-J**) Electron micrographs of longitudinal sections through single CEP sense organs in adult (E) wild-type male, (F) male with feminized glia, and (G) *mab-3(mu15)* mutant male, (H) wild-type hermaphrodite, (I) hermaphrodite with masculinized glia, and (J) *nfya-1(ok1174)* mutant hermaphrodite. First annulus (bracket) provides a landmark to locate the normal position of the pore. Arrows, cuticle pore; arrowheads, extracellular vesicles. Scale bars, 1 µm. The fraction of CEP sense organs exhibiting the represented phenotype out of the total number scored is shown. See also Figure S5.

Next, to directly test whether male-specific CEPso glial gene expression is required for cuticle pore formation, we used electron microscopy to visualize the aECM overlying the CEP sense organs of wild-type males and males that fail to express male-specific genes in their glia. We obtained longitudinal serial sections through the heads of wild-type males, males with feminized glia, and *mab-3* mutant males at the adult stage (∼40-100 serial sections per animal, 3-4 animals per genotype; see Figure S5C). CEM neurons were identified based on their position relative to the mechanosensory CEP neuron, their morphology, and the presence of extracellular vesicles in the space surrounding the CEM endings. In wild-type males, we observe that each CEM cilium protrudes through a pore that is consistently located anterior to the first annulus (circumferential ridge (Page, 2007)) of the cuticle (Figure 5E, n=9/9). In males with feminized glia, the CEM cilia are misdirected inward, consistent with the fluorescent images, and pores are absent in most of the sense organs analyzed (Figure 5F, n=6/7). Similarly, in *mab-3* mutant males, CEM cilia are misdirected inward and pores are absent (Figure 5G, n=10/10). We also observe that there is an overaccumulation of extracellular vesicles (EVs) in males with feminized glia and *mab-3* mutant males, suggesting that the EVs are trapped inside the animal and cannot be released into the external environment (Wang et al., 2014). Thus, male-specific gene expression in CEPso glia is necessary for cuticle pore formation.

To determine whether male-specific glial gene expression in CEPso glia is sufficient to remodel the aECM into an open pore, we obtained longitudinal serial sections of adult wild-type hermaphrodites and hermaphrodites that inappropriately express male genes (∼30-100 serial sections per animal, 3-4 animals per genotype; see Figure S5D). Since hermaphrodites do not have CEM neurons, the mechanosensory CEP neuron was used as a landmark to locate the relevant serial sections to analyze (Figure 5H). Strikingly, hermaphrodites with masculinized glia often have ectopic cuticle pores positioned anterior to the first annulus (Figure 5I, n=7/11). Ectopic pores are also present in *nfya-1* mutant hermaphrodites (Figure 5J, n=13/13). The pores are properly patterned despite the absence of the CEM neuron. This shows that forced expression of male-specific genes in CEPso glia is sufficient for driving cuticle pore formation. Together, these findings reveal that a sex-specific switch in glial gene expression remodels the aECM from a closed sheet to an open pore (Figures S5C-S5D).

## DISCUSSION

A long-standing framework for understanding the formation of biological structures distinguishes between “self-assembly,” where components come together without energy input to attain a stable, thermodynamically-favored conformation, and “self-organization,” where energy input is continuously required to actively maintain a non-equilibrium state (Marshall, 2020; Misteli, 2001). However, a different approach may be needed to understand patterning of aECM structures. Although most aECM structures are highly stable – persisting even after death of the organism – previous studies show that the initial formation of these structures often requires extensive energy input and cellular activity. For example, in *Drosophila,* epidermal cells and olfactory hair cells form tooth-like denticles and peg-like extensions, respectively, that act as molds to sculpt newly secreted matrix proteins, and then retract as the aECM solidifies (Ando et al., 2019; Fernandes et al., 2010). Actin networks also shape aECM pattern through contraction of the cell surface and localized secretion, as in alae of the *C. elegans* cuticle and taenidia of the *Drosophila* trachea (Katz et al., 2022; Öztürk-Çolak et al., 2016b). Finally, cellular movements apply mechanical forces that can stretch or fold the aECM during tube morphogenesis (Dong and Hayashi, 2015; Gill et al., 2016). In these examples, aECM structures are thought to be sculpted mechanically by active movements of the underlying cells. By contrast, our findings suggest a different paradigm in which secreted matrix molecules can “self-assemble” into a closed sheet or open pore conformation without major cellular rearrangements. Understanding how aECM can be remodeled through changes in its biochemical composition will lead to new strategies for manipulating matrix structure *in vitro* and *in vivo*. Applications may include tissue engineering, improving drug delivery across epithelia, and treatment of aECM disorders such as cystic fibrosis or chronic obstructive pulmonary disorder.

Our work is consistent with recent evidence that a transient, or provisional, matrix initially patterns the mature aECM and is then removed (Cohen et al., 2020; Gill et al., 2016; Katz et al., 2022; Öztürk-Çolak et al., 2016a). We propose that GRL-18 is a transient matrix component that forms temporary rings at sites of future cuticle pores, where it may act as part of a physical plug to keep the cuticle open or as a corral to retain pore-forming components in a restricted region. Unsurprisingly, GRL-18 is not the sole determinant of pore formation – by examining *grl-18* mutant males, we find that the CEM cilia of *grl-18* mutant males bend outwards normally, suggesting that pores are still formed (Figures S5A-S5B). Rather, GRL-18 is likely to be part of a battery of aECM components that are coordinately expressed in male CEPso glia to induce pore formation; for example, we find that a transcriptional reporter for the collagen-encoding gene *col-53* is also expressed constitutively in ILso glia and switches on in CEPso glia only at sexual maturity in males, like *grl-18*. Many Grl domain proteins and other Hedgehog-related proteins (Grd, Wrt, Qua domains) of *C. elegans* are expressed specifically in various socket glia or other cuticle-producing cells (Hao et al., 2006). It is intriguing to consider that this protein family may have a shared role in patterning aECM rather than acting as diffusible signaling molecules – indeed, perhaps Hedgehog proteins themselves evolved from ancient structural components of the aECM.

Finally, we discovered a novel sexual dimorphism in glia and identified some of its regulators. Sex differences have been extensively studied in neurons, but little is known about how these differences are controlled in glia. Glial sex differences are of high interest in the mammalian brain, because glia are a major cell type in the nervous system that sculpts neuronal connections and modulates neuronal activity. Adult male and female microglia have distinct gene expression profiles and can influence sex-specific neuronal structure and function (Guneykaya et al., 2018; Lenz et al., 2013; Villa et al., 2018); astrocyte glia exhibit molecular differences early in development and differ in number and morphology in brain regions involved in pheromone-sensing and hormone secretion (Amateau and McCarthy, 2002; Rurak et al., 2022; Schwarz and Bilbo, 2012); and glial functions are disrupted in disease models for sex-biased disorders, including autism and Alzheimer’s disease (Neniskyte and Gross, 2017; Vegeto et al., 2020). In most cases, it remains unclear whether glial sex differences are a response to a sex-specific neuronal environment or reflect cell-intrinsic differences in the glia themselves, and what regulators might control these differences. In *C. elegans*, two examples have been described in which glia play sex-specific roles as neuronal progenitors (Molina-García et al., 2020; Sammut et al., 2015). We show that a sex-specific switch can also alter function of the mature glial cell itself. This switch is controlled cell autonomously by some of the same regulators that control sex differences in neurons (sex identity: *fem-3*, *tra-2*, *mab-3*; timing: *lep-2*, *lep-5*) but with novel downstream components (*nfya-1*, *bed-3*, *jmjd-3.1*) that may be more glial-specific.

Altogether, our findings underscore that aECM is not a static barrier but can contain highly patterned local structures that are constructed through coordinated regulation of discrete gene expression modules. In this sense, the aECM identity of a cell can be viewed as a defining feature of a cell type, much like neurotransmitter identity is for neurons.

## METHODS

### Strains and plasmids

All strains and plasmids in this study are in Tables S2 and S3. All strains are in an N2 background and, unless otherwise stated, contain *him-5(e1490)* or *him-8(e1489)* to generate a higher percentage of male progeny. Animals were grown at 20-22°C on nematode growth media (NGM) plates seeded with *E. coli* OP50 bacteria (Brenner, 1974). Transgenic strains were generated using standard techniques (Mello and Fire, 1995) with injections of 100 ng/μL DNA (5-80 ng/μL per plasmid). Unless otherwise stated, all animals were picked to sex-segregated plates as L4 hermaphrodites and males based on vulva and tail morphology, respectively, and scored as 1-day adults the next day.

### Generation of alleles by genome editing

To generate the endogenous *grl-18* transcriptional reporter (*grl-18-*SL2-YFP:H2B), a SapTrap plasmid (Dickinson et al., 2015; Schwartz and Jorgensen, 2016) was assembled to contain the gRNA (5’GAGTATCAAAGTTTGAATCA-3’) and repair template. The repair template includes ∼530 bp 5’ and 3’ homology arms flanking an SL2-YFP:H2B fragment, a loxP-flanked positive selection marker, *sqt-1*, that causes a roller (Rol) phenotype, and a sequence encoding heat-shock-inducible *Cre* recombinase. A total of 65 ng/μL SapTrap plasmid was co-injected with a panel of markers used to select against extrachromosomal arrays (2 ng/μL pCFJ90, 4 ng/μL pGH8, 4 ng/μL pCFJ104, and 25 ng/μL pBlueScript) (Dickinson et al., 2015) into the strain EG9888, which carries an integrated transgene for germline Cas9 expression (Schwartz et al., 2021). A single genome-edited line was isolated exhibiting the Rol phenotype and confirmed by Sanger sequencing to contain the SL2-YFP:H2B fragment inserted after the stop codon of the endogenous *grl-18* locus (Table S4). Finally, the integrated Cas9 transgene was removed by outcrossing to wild-type animals and the positive-selection marker was removed by Cre induction via heat-shock (34°C for 4 h).

To generate a null allele of *grl-18*, a co-CRISPR (Arribere et al., 2014; Kim et al., 2014) and dual-sgRNA (Chen et al., 2014) approach was used. Two gRNAs were selected to target the 5’ and 3’ ends of the *grl-18* gene (5’-AGTTTACCGAATCCAAGTTG-3’ and 5’GAGTATCAA AGTTTGAATCA-3’, respectively) and cloned into separate plasmids. To screen and select for genome-edited animals, *unc-58* was targeted to introduce a gain-of-function allele corresponding to *unc-58(e665gf)* (Arribere et al., 2014). 25 ng/μL of each sgRNA plasmid was co-injected with 25 ng/μL pJA50 (*unc-58* sgRNA plasmid), 600 ng/μL *unc-58(e665)* repair oligo, and 25 ng/μL pBlueScript into the strain EG9888. Animals carrying a deletion in *grl-18* were identified via PCR using primers flanking the gRNA target sites. The *hmn341* allele was recovered as a 960 bp deletion that removes the conserved Grl domain sequences of the endogenous *grl-18* locus (Table S4).

The allele *grl-18(syb6299)* was generated by SunyBiotech (Fuzhou, China). The first intron of the endogenous *grl-18* locus was deleted and sfGFP was inserted between the first and second exon. Synonymous mutations were introduced to prevent re-cutting by Cas9/gRNA (Table S4).

### Forward genetic screens for altered CEPso glial gene expression

Novel alleles in Table 1 were isolated through two clonal screens (*hmn289*, *hmn304*, and *hmn305*) and one nonclonal screen (*hmn316*, *hmn317*, *hmn318*, and *hmn319*). In all cases, animals were mutagenized with 70 mM ethyl methanesulfonate (EMS, Sigma) at 22°C for 4 h. In each of two independent clonal screens, animals of genotype *hmnIs47* (*grl-18*pro:mApple) I; *him-5* V were mutagenized, ∼600 F1 progeny were picked to individual plates, and plates of F2 siblings were screened using a fluorescence stereomicroscope to identify the presence of adult males lacking *grl-18*pro:mApple expression in CEPso glia. When such males were found, the mutant allele was recovered by picking ∼6-12 L4 hermaphrodite siblings to individual plates. Siblings that produced male progeny lacking *grl-18*pro:mApple in CEPso glia carried the mutant allele of interest. In a single nonclonal screen, animals of genotype *hmnIs82* (*grl-18*pro:GFP) II were mutagenized and pooled F2 progeny were screened *en masse* using a fluorescence stereomicroscope to identify and isolate adult hermaphrodites that inappropriately express *grl-18*pro:GFP in CEPso glia. Mutant alleles generated in this study are in Table S4.

### Genetic mapping and identification of causal mutations

Causal mutations in *hmn289* and *hmn305* were identified by candidate gene analysis based on distinctive tail morphology defects observed in mutant males. *hmn289* males exhibit missing tail sensory rays, a phenotype that is characteristic of *mab* (*m*ale *ab*normal) mutants (Hodgkin, 1983), and showed non-complementation with *mab-3(mu15)* (from the cross *mu15*/+ x *hmn289*, 34/60 1-day adult males failed to express *grl-18*pro:mApple in CEPso glia), which is known to affect male-specific phenotypes (Fagan et al., 2018; Mason et al., 2008; Ross et al., 2005; Shen and Hodgkin, 1988; Yi et al., 2000). *hmn305* males exhibit a *lep* (*lep*toderan) male tail phenotype, characterized by retention of the juvenile tail tip in the adult stage (Nguyen et al., 1999), and showed non-complementation with *lep-2(ok900)* (from the cross *ok900*/+ x *hmn305*, 26/53 1-day adult males failed to express *grl-18*pro:mApple in CEPso glia), which is known to affect the timing of sexual maturation (Herrera et al., 2016; Lawson et al., 2019). Sanger sequencing showed that *hmn289* encodes a R90C missense mutation in the second DNA-binding domain of MAB-3 and that *hmn305* encodes a C216Y missense mutation in the conserved RING finger domain of LEP-2. The reference alleles *mab-3(mu15)* and *lep-2(ok900)* recapitulated the *grl-18* reporter defects of *hmn289* and *hmn305* (Table 1). Together, these data indicate that *hmn289* and *hmn305* are alleles of *mab-3* and *lep-2*, respectively.

To identify the causal mutation in *hmn304*, a series of crosses were first used to determine that the mutation is X-linked. Whole-genome sequencing identified a W831STOP nonsense mutation in *jmjd-3.1* that is predicted to truncate the protein within the conserved JmjC DNA-binding domain. The *jmjd-3.1(gk387)* mutation recapitulates the *hmn304* phenotype (Table 1) and a transgene bearing the fosmid WRM0610dB04, which consists of the 4.4 kb *jmjd-3.1* gene with ∼15 kb upstream and ∼13 kb downstream sequences, rescues the *hmn304* mutant phenotype (41/42 1-day adult males express *grl-18*pro:GFP in at least one CEPso glia). Together, these data indicate that *hmn304* is an allele of *jmjd-3.1*.

To identify the causal mutation in *hmn318*, whole-genome sequencing was performed. Among the mutations in this strain, we identified a C144G missense mutation in the DNA-binding BED domain of BED-3, a transcriptional repressor important in vulval development (Inoue and Sternberg, 2010). We noted that *hmn318* hermaphrodites exhibit weak vulval defects, consistent with loss of *bed-3* function. Further, we found that two existing alleles of *bed-3*, *sy705* and *gk996*, recapitulate the *hmn318* defects in *grl-18*pro:GFP expression (Table 1). A transgene bearing the fosmid WRM0624bB06, which consists of the 5.4 kb *bed-3* gene with ∼19 kb upstream and ∼7 kb downstream sequences, rescues the *hmn318* mutant phenotype (9/67 1-day adult hermaphrodites inappropriately express *grl-18*pro:GFP in at least one CEPso glia). Together, these data indicate that *hmn318* is an allele of *bed-3*.

To identify the causal mutations in *hmn316*, *hmn317*, and *hmn319*, one-step mapping was performed by crossing each mutant to the polymorphic strain CB4856 (“Hawaiian”), selecting F2 recombinants that exhibit inappropriate *grl-18* expression in CEPso glia, and analyzing their pooled progeny by whole-genome sequencing (Doitsidou et al., 2010). Sequencing results were analyzed using MiModD (v0.1.9) (Maier, 2018) through a local Galaxy interface (Afgan et al., 2018). All three mutations were mapped to an interval between 6 Mb and 14 Mb on chromosome X. Each mutant was found to have probable loss-of-function sequence changes in *nfya-1*: *hmn316*, g>a splice site donor mutation between exons 4 and 5; *hmn317*, R21STOP nonsense mutation; and *hmn319*, Q146STOP nonsense mutation. A strain bearing the reference allele *nfya-1(ok1174)* recapitulated the *grl-18* reporter expression defect of all three mutants (Table 1). In complementation tests, *hmn316* failed to complement *hmn317*, *hmn319*, or *ok1174* (41/42, 27/27, and 26/26 1-day adult hermaphrodites inappropriately express *grl-18*pro:GFP in CEPso glia, respectively). Finally, the *hmn316* mutant phenotype was rescued with the WRM0637aE09 fosmid, which consists of the 2.4 kb *nfya-1* gene with ∼14 kb upstream and ∼17 kb downstream sequences (5/76 1-day adult hermaphrodites inappropriately express *grl-18*pro:GFP in CEPso glia), and the *ok1174* mutant phenotype was rescued with *nfya-1* cDNA (Figure 3C). Together, these results indicate that *hmn316*, *hmn317*, and *hmn319* are alleles of *nfya-1*.

### Fluorescence microscopy and image processing

Animals were washed and immobilized in M9 solution containing 50 mM sodium azide and mounted on 2% agarose pads containing 50 mM sodium azide. Image stacks were collected on a DeltaVision Core imaging system (Applied Precision) with a UApo 40x/1.35 NA, PlanApo 60x/1.42 NA, or UPlanSApo 100x/1.40 NA oil immersion objective and a CoolSnap HQ2 camera. Images were deconvolved using Softworx (Applied Precision) and maximum intensity projections were generated in ImageJ (Fiji) (Schindelin et al., 2012). The brightness and contrast of each projection were linearly adjusted in Affinity Photo 1.10.1. Fluorescent signals were pseudo-colored, and merged images were generated using the Screen layer mode in Affinity Photo 1.10.1.

### Developmental staging for time course experiments

For time course experiments, single animals were tracked across development from larval to adult stages. L3 hermaphrodites and males were identified based on vulva and tail morphology, respectively, picked to individual plates, and scored. The same individuals were scored after ∼10 h (late L4), again after ∼12 h (1-day adult), and then at 24 h intervals (2-, 3-, 4-day adults). Animals that crawled off the plate or died before the experiment was complete were not included in the data. Fluorescent reporter expression was scored visually on a Nikon SMZ1500 stereomicroscope with an HR Plan Apo 1.6x objective.

To image animals during the L4/adult molt, L4 males were selected based on tail morphology and either imaged immediately or monitored up to 4 h until the cuticle was shed and the adult tail was formed. Animals were imaged as described above (“Fluorescence microscopy and image processing”). Staging within the molt was determined based on male tail morphology (Emmons, 2005; Nguyen et al., 1999) as follows: L4 – pointed tail, no retraction of hypodermal cells; early molt – rounded tail, hypodermal cells hyp9 and hyp10 have retracted from the L4 cuticle; late molt – further retraction of hypodermal cells hyp8-11 from the L4 cuticle, rays beginning to form; adult – L4 cuticle has been shed, fans and rays are fully formed.

### Binary scoring of CEM cilia trajectories

CEM and CEP cilia were imaged using fluorescent markers for each neuron (CEM, *pkd-2*pro:GFP; CEP, *dat-1*pro:mApple). L4 animals were selected based on tail morphology and imaged as 1-day adults as described above (“Fluorescence microscopy and image processing”). A sense organ was only scored if both CEM and CEP were labeled and oriented flat with their cilia in the x-y plane, not the z plane. In rare cases, either one or both the CEM and CEP cilia were short or abnormally shaped; these were not scored. The CEP cilium was used as a reference for a trajectory that bends inward. If the CEM cilium traveled in the same direction as the CEP cilium, then it was scored as “bent in”; if it traveled in the direction orthogonal to the CEP cilium, then it was scored as “bent out”. Because the CEM cilium is extremely thin and the fluorescent signal at the tip is very dim, the brightness and contrast of the images were adjusted until the cilia were visible, which often saturated the signal in the rest of the CEM ending.

### Electron microscopy

Samples were high-pressure frozen and quick freeze substituted as previously described (Kolotuev, 2014). Epon flat-embedded samples were carefully oriented for longitudinal sectioning through the head, except for a wild-type male and hermaphrodite that were each oriented for transverse sectioning (Burel et al., 2018; Kolotuev, 2014). To target the region of interest and to minimize artifacts, excess resin was removed using a 90° trimming tool (Diatome, Switzerland). Samples were sectioned using an ATS knife (Diatome, Switzerland) mounted on a Leica UC7 ultramicrotome (Leica, Austria). Approximately 100 to 300 sections were obtained per animal and transferred to a 2x4 cm silicon wafer (Burel et al., 2018; Franke and Kolotuev, 2021). Wafers were dried at ambient temperature by evaporation and subsequently incubated in a 60°C oven for further fixation as previously described (Burel et al., 2018; Franke and Kolotuev, 2021; Santella et al., 2022). Wafers were analyzed with a Helios SEM microscope (Thermo Fisher Scientific) at 2keV landing energy and 0.8 nA beam current at 2 mm distance using a Mirror Detector (MD-BSA) (Burel et al., 2018; Franke and Kolotuev, 2021). Images were collected manually or automatically with 4-6 µs dwell time using Maps 3.11 software (Thermo Fisher Scientific) (Burel et al., 2018; Franke and Kolotuev, 2021; Santella et al., 2022). Images were collected at 3 mm/s (1024x886) or 5 mm/s (6084x2044) to generate 20 nm and 5 nm resolution images, respectively. To cover a larger area of the sample, several images were collected at a given resolution and stitched together using Maps 3.17 software (Thermo Fisher Scientific) (Burel et al., 2018; Franke and Kolotuev, 2021). Finally, serial sections were aligned using the IMOD program (University of Colorado) (Kremer et al., 1996).

## ACKNOWLEDGMENTS

We thank Douglas Portman, Maureen Barr, and members of the Scott Kennedy laboratory for reagents; Lisa Goodrich, Joshua Kaplan, Constance Cepko, and Norbert Perrimon for advice; members of the Heiman laboratory for comments on the manuscript; and Wormbase. Some strains were provided by the *C. elegans* Genetics Center [CGC, funded by National Institutes of Health Office of Research Infrastructure Program (P40 OD010440)]. This work was supported by NIH grant R01NS124879 to M.G.H. and by NIH grant F31NS122139 to W.F.

## Notes

### Competing Interest Statement

The authors have declared no competing interest.

