## Supplemental Material for "A sex-specific switch in a single glial cell patterns the apical extracellular matrix"

**Supplmental Figure S1 - S5**
**Supplmental Tables S1 - S4**

SUPPLEMENTAL FIGURE S1

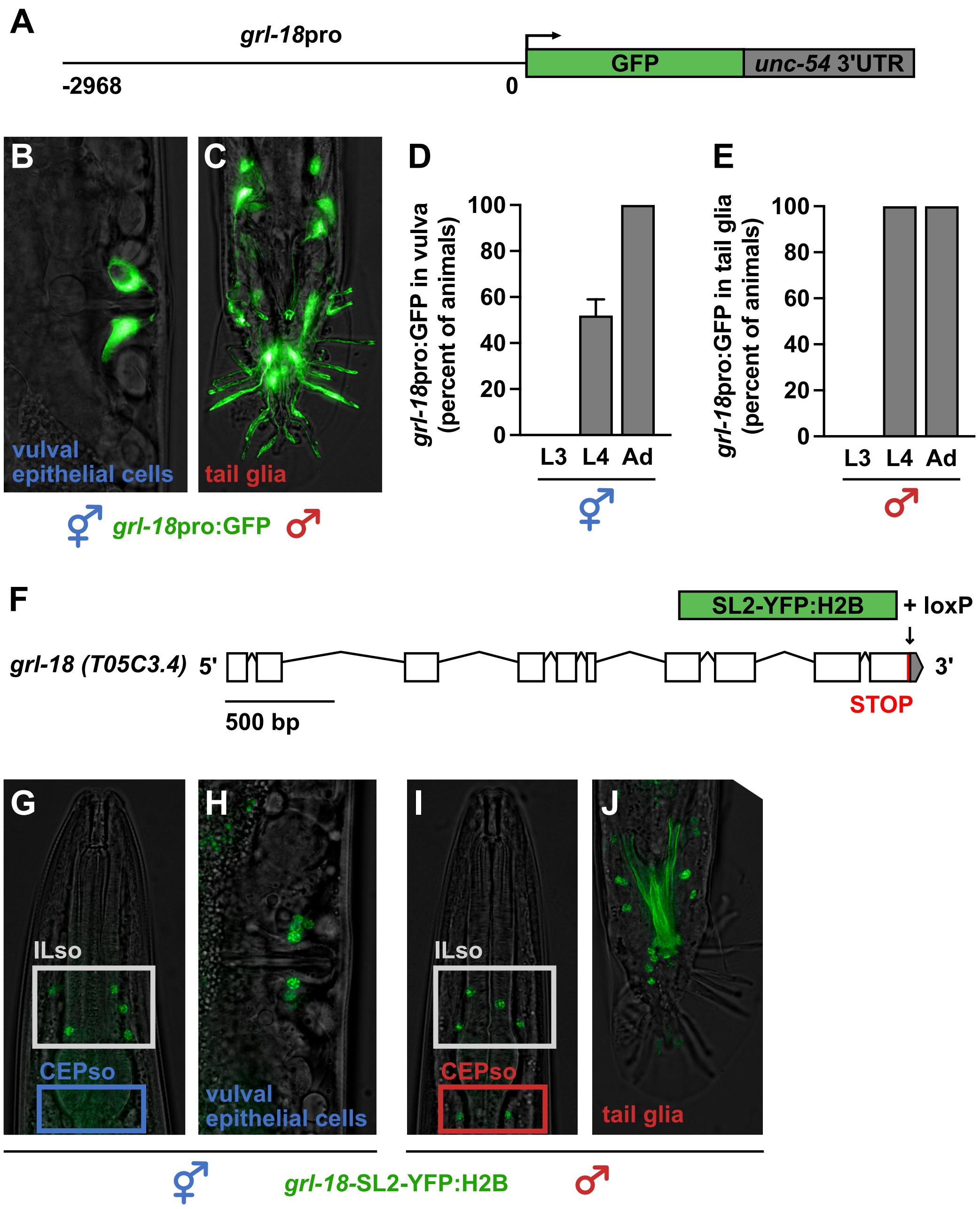


**Supp. Fig. S1. Transgenic and endogenous *grl-18* reporters show sex-specific expression.**

(**A**) Schematic of transcriptional reporter transgene *grl-18*pro:GFP, containing 2968 bp upstream of the translational start site. (**B-C**) Expression of *grl-18*pro:GFP in 1-day adult (B) hermaphrodite vulval epithelial cells and (C) male tail glia, including ray structural and hook socket glia. (**D-E**) Fraction of hermaphrodites (n=48) or males (n=50) at the indicated ages showing *grl-18*pro:GFP expression in (D) vulval cells and (E) tail glia. The same individuals were followed and scored as in Figure 1D and 1E. Error bars, SEM. (**F**) Schematic of *grl-18* endogenous transcriptional reporter. White boxes, exons; lines, introns; gray, 3' untranslated region (UTR). An insertion encoding the splice leader 2 (SL2) sequence followed by a YFP-tagged histone H2B (YFP:H2B) was inserted immediately after the stop codon. A loxP site and short linker sequence remain after removal of a selection cassette. (**G-J**) Expression of *grl-18* endogenous reporter in 1-day adult hermaphrodite (G) ILso nuclei and (H) four pairs of nuclei in the vulva and in 1-day adult male (I) ILso, CEPso, and (J) tail glia nuclei. Due to relatively dim expression, autofluorescence is visible in spicules of the mail tail.

### SUPPLEMENTAL FIGURE S2

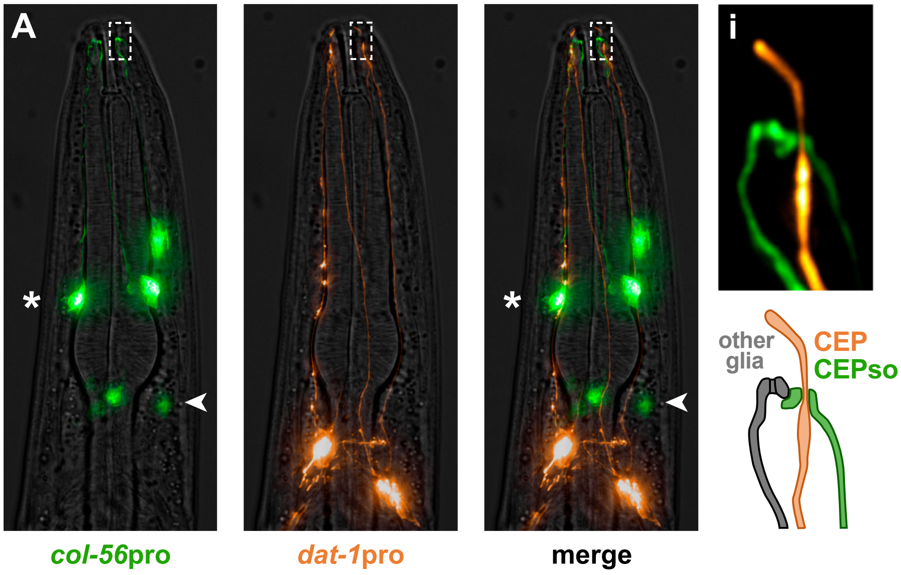


**Supp. Fig. S2. Identification of a novel marker for CEPso glia.**

(**A**) *col-56* was identified as a novel marker for CEPso glia (arrowhead), which were identified by their cell body position posterior to the first pharyngeal bulb and by their endings that form a channel wrapping the CEP cilium (inset, **i**). *col-56* is also expressed in OLso glia (asterisk). Green, *col-56*pro:GFP in CEPso and OLso glia; orange, *dat-1*pro:mApple in CEP neurons.

SUPPLEMENTAL FIGURE S3

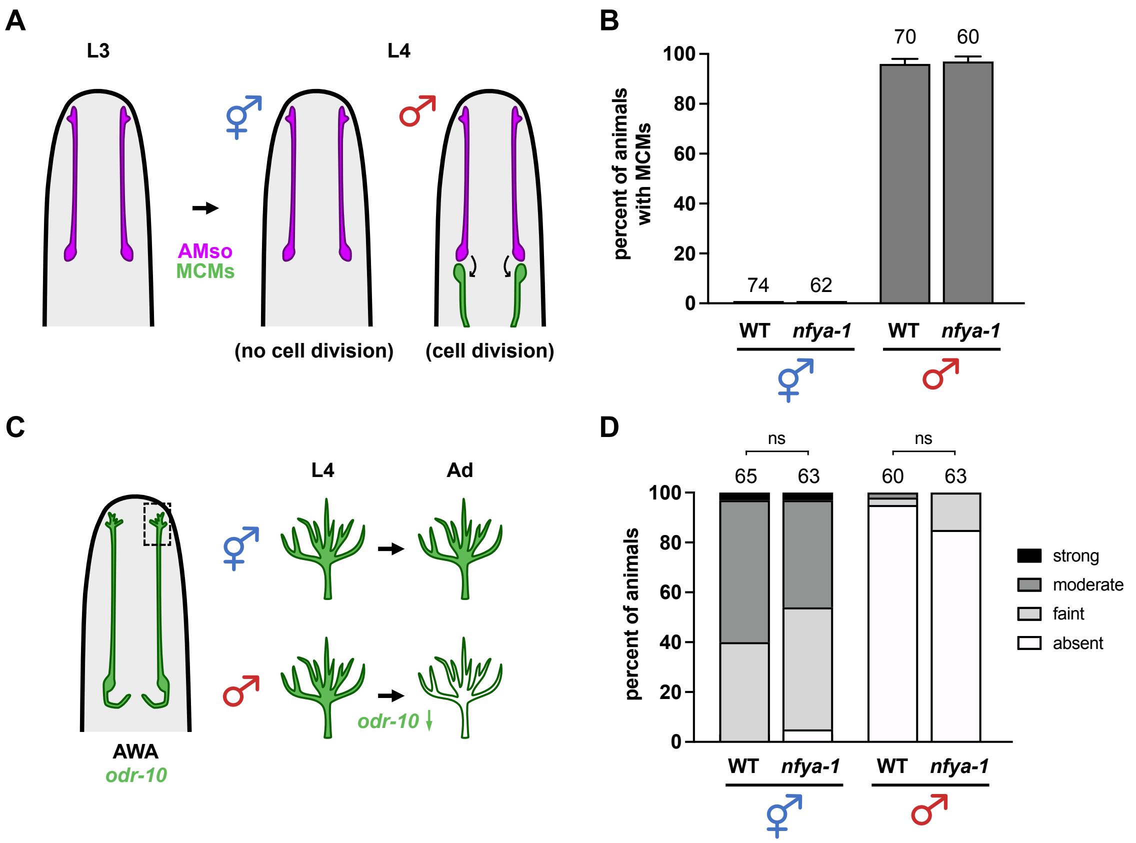


**Supp. Fig. S3. *nfya-1* does not affect other male-specific phenotypes in glia and neurons.**

(**A**) Schematic showing amphid socket glia (AMso, magenta) divide to generate MCM interneurons (green) at the L4 stage only in males (Sammut et al., 2015). (**B**) Fraction of 1-day adult wild-type or *nfya-1(ok1174)* mutant animals of each sex with MCM interneurons, as visualized by *pdf-1*pro:RFP. Sample sizes are indicated above the bars. Error bars, SEM. (**C**) Schematic showing that expression of the ODR-10 chemoreceptor (ODR-10:GFP) in AWA neuronal cilia is reduced or absent in adult males compared to adult hermaphrodites (Ryan et al., 2014). (**D**) Fraction of 1-day adult wild-type or *nfya-1(ok1174)* mutant animals of each sex with ODR-10:GFP expression at the indicated levels. Expression levels were binned as 0, absent; 1, faint; 2, moderate; and 3, strong, and the Mann-Whitney test was used as previously described (Lawson et al., 2019; Ryan et al., 2014; Wexler et al., 2020); ns = no statistically significant difference ($p > 0.05$).

SUPPLEMENTAL FIGURE S4

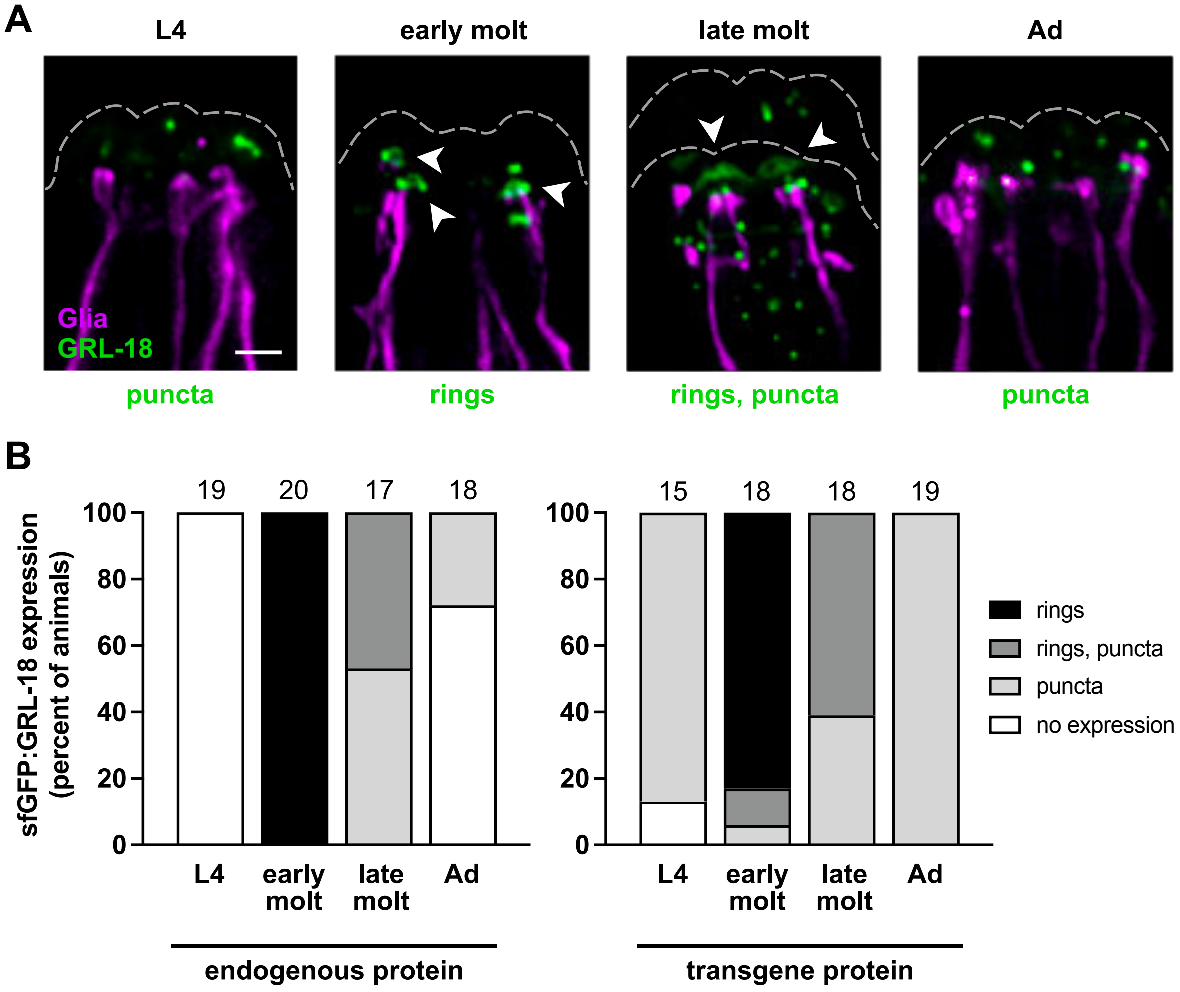


**Supp. Fig. S4. sfGFP-tagged GRL-18 localizes to transient rings near glial endings.**

(**A**) Consistent with results obtained using an endogenous insertion, sfGFP-GRL-18 expressed from a low-copy transgene localizes to rings at the nose tip in males during the L4 to adult (Ad) cuticle molt; sfGFP tag is inserted at the N-terminus of full-length GRL-18 protein (see Figure 4F). Arrowheads, sfGFP-tagged GRL-18 rings. Scale bar, 2 μm. **(B)** The fraction of males at the indicated stages that show no expression or localization to rings, diffuse puncta, or both is similar between strains with an endogenous insertion (left) or low-copy transgene (right). Sample sizes are indicated above the bars.

### SUPPLEMENTAL FIGURE S5

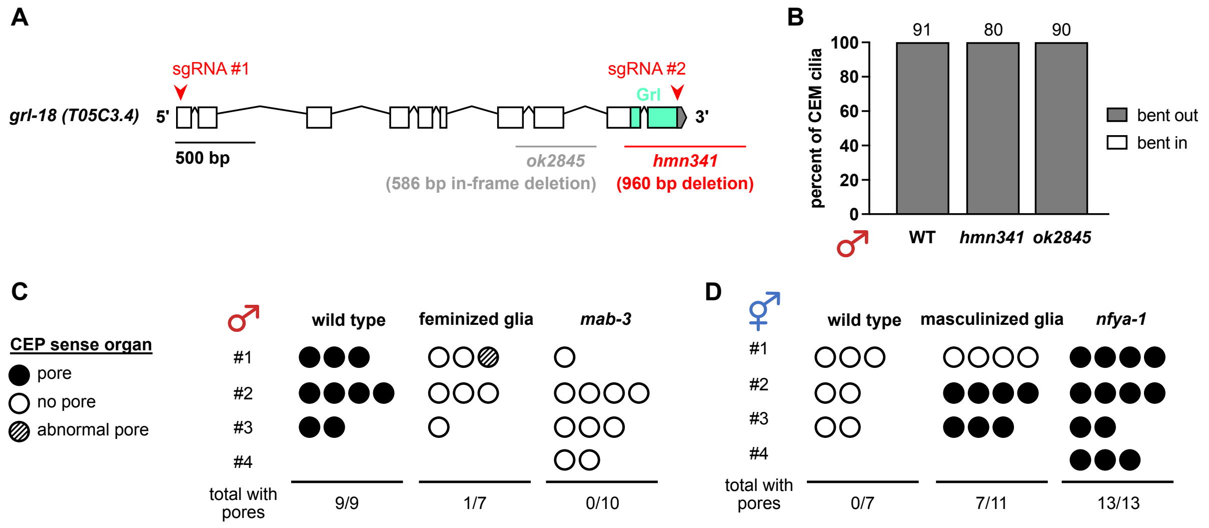


**Supp. Fig. S5. Male-specific gene expression in CEPso glia causes cuticle pore formation.**

(**A**) Schematic of the *grl-18* gene showing existing allele *ok2845* and a novel allele *hmn341*. *ok2845* is an in-frame deletion that is predicted to spare the Grl domain. *hmn341* deletes the sequence encoding the Grl domain. White boxes, exons; lines, introns; gray, 3' UTR. Red arrowheads indicate sites targeted in the dual-sgRNA CRISPR/Cas9 approach used to generate the *hmn341* deletion. (**B**) Quantification of CEM cilia trajectories in *grl-18* mutants. Sample sizes are indicated above the bars. **(C, D)** Summary of observations made for each CEP sense organ scored in EM serial sections of adult (C) males and (D) hermaphrodites (n=3-4 animals per genotype, numbered #1-#4). Each animal has four CEP sense organs, but due to missing or damaged sections, not all four were scored in every animal. Filled circle, cuticle pore; open circle, no pore; striped circle, abnormal pore. In the sense organ with an abnormal pore, the CEM ending appears to break through the cuticle adjacent to the CEP nubbin, reminiscent of a phenotype previously described for CEP in *cat-6* mutants (Perkins et al., 1986).

### SUPPLEMENTAL TABLE S1

| | Genotype | Age | *grl-18*$^{+}$ CEPso glia (percent of animals) | n |
|---|---|---|---|---|
| ♂ | I. DM Domain Transcription Factors | | | |
| | wild type males | 1d | 100 | 67 |
| | *mab-23* | 1d | 98[a] | 58 |
| | *dmd-3* | 1d | 100 | 68 |
| | *dmd-5* | 1d | 98[a] | 42 |
| | *dmd-6* | 1d | 100 | 61 |
| | *dmd-7* | 1d | 100 | 69 |
| | *dmd-8* | 1d | 100 | 41 |
| | *dmd-9* | 1d | 100 | 68 |
| | *dmd-10* | 1d | 97[a] | 71 |

[a] phenotype is partially penetrant; at least one CEPso fails to express marker

**Supp. Table S1. Other DM domain transcription factors are not required for sex-specific CEPso glial gene expression.**

Percent of 1-day adult (1d) males expressing the transcriptional reporter *grl-18*pro:mApple in CEPso glia for mutants in DM domain transcription factors other than *mab-3* (Table 1), except the lethal mutant *dmd-4*. Expression in ILso glia and male tail glia were unaffected.

### SUPPLEMENTAL TABLE S2

### Supp. Table S2. Strains used in this study.

### (A) Strains from previous studies

| Strain | Genotype | Figures, Tables | References |
|---|---|---|---|
| CHB3829 | *hmnIs82* [*grl-18*pro:GFP] II | 1B-1E, S1B-S1E | Fung et al., 2020 |

### (B) Strains generated for this study

| Strain | Genotype | Figures, Tables |
|---|---|---|
| CHB3843 | *hmnIs82* [*grl-18*pro:GFP] II; *him-5(e1490)* V | 2B, 2F, 3C-3D, 5E, 5H, Table 1, Table S5C-S5D |
| CHB3925 | *hmnIs82* [*grl-18*pro:GFP] II; *him-5(e1490)* V; *hmnEx2172* [*prab-3*:*fem-3*-mCherry] | 2B |
| CHB3900 | *hmnIs82* [*grl-18*pro:GFP] II; *him-5(e1490)* V; *hmnEx2193* [*mir-228*pro:*fem-3*; *unc-122*pro:RFP] | 2B |
| CHB4032 | *hmnIs82* [*grl-18*pro:GFP] II; *him-5(e1490)* V; *hmnEx2215* [*col-56*pro:*fem-3*; *unc-122*pro:RFP] | 2B |
| CHB4298 | *nIs131* [*pkd-2*:GFP] I; *hmnIs82* [*grl-18*pro:GFP] II; *him-5(e1490)* V | 2D, 2H |
| CHB4285 | *nIs131* [*pkd-2*:GFP] I; *hmnIs82* [*grl-18*pro:GFP] II; *him-5(e1490)* V; *ceh-30(n3714)* X | 2D |
| CHB4030 | *hmnIs82* [*grl-18*pro:GFP] II; *him-5(e1490)* V; *hmnEx2208* [*prab-3*:*tra-2*-mCherry] | 2F |
| CHB3926 | *hmnIs82* [*grl-18*pro:GFP] II; *him-5(e1490)* V; *hmnEx2204* [*mir-228*pro:*tra-2*; *unc-122*pro:GFP] | 2F |
| CHB4031 | *hmnIs82* [*grl-18*pro:GFP] II; *him-5(e1490)* V; *hmnEx2211* [*col-56*pro:*tra-2*; *unc-122*pro:RFP] | 2F |
| CHB4338 | *nIs131* [*pkd-2*:GFP] I; *hmnIs82* [*grl-18*pro:GFP] II; *him-5(e1490)* V; *ceh-30(n4289)* X | 2H |
| CHB4185 | *hmnIs82* [*grl-18*pro:GFP] II; *him-5(e1490)* V; *nfya-1(ok1174)* X | 3C-3D, 5J, Table 1, Table S5D |
| CHB4245 | *hmnIs82* [*grl-18*pro:GFP] II; *him-5(e1490)* V; *nfya-1(ok1174)* X; *hmnEx4304* [*nfya-1*pro:*nfya-1*-cDNA; *unc-122*pro:RFP] | 3C |
| CHB4303 | *hmnIs82* [*grl-18*pro:GFP] II; *him-5(e1490)* V; *nfya-1(ok1174)* X; *hmnEx2329* [*col-56*pro:*nfya-1*-cDNA; *unc-122*pro:RFP] | 3C |
| CHB4281 | *nfyb-1(cu13)* II; *him-5(e1490)* V; *hmnIs78* [*grl-18*pro:GFP] X | 3C Table 1 |
| CHB4603 | *nfyc-1(tm4541)* II; *him-5(e1490)* V; *hmnIs78* [*grl-18*pro:GFP] X | 3C, Table 1 |
| CHB3845 | *hmnIs82* [*grl-18*pro:GFP] *mab-3(mu15)* II; *him-5(e1490)* V | 3D, 5G, Table S5C |
| CHB4220 | *hmnIs82* [*grl-18*pro:GFP] *mab-3(mu15)* II; *him-5(e1490)* V; *nfya-1(ok1174)* X | 3D |
| CHB4641 | *hmnIs47* [*grl-18*pro:mApple] I; *him-8(e1489)* IV; *grl-18(syb6299*[*sfGFP:grl-18*]*)* V | 4A-4C, S4B |
| CHB4467 | *hmnIs47* [*grl-18*pro:mApple] I; *him-8(e1489)* IV; hmnEx2400 [*grl-18*pro:sfGFP:*grl-18*-gDNA; *rol-6(su1006)*] | 4G, S4A-S4B |
| CHB4642 | *hmnIs47* [*grl-18*pro:mApple] I; *him-8(e1489)* IV; *hmnEx2486* [*grl-18*pro:*grl-18*-gDNA:sfGFP; *rol-6(su1006)*] | 4H |
| CHB4465 | *hmnIs47* [*grl-18*pro:mApple] I; *him-8(e1489)* IV; *hmnEx2398* [*grl-18*pro:sfGFP:*grl-18*-gDNA(352-445); *rol-6(su1006)*] | 4I |
| CHB4461 | *hmnIs47* [*grl-18*pro:mApple] I; *him-8(e1489)* IV; *hmnEx2394* [*grl-18*pro:sfGFP:*grl-18*-gDNA(31-351); *rol-6(su1006)*] | 4J |

| | | |
|---|---|---|
| CHB4519 | *him-5(e1490)* V; *hmnEx2422* [*pkd-2*pro:GFP; *dat-1*pro:mApple; *rol-6(su1006)*] | 5B, 5D, S5B |
| CHB4543 | *him-5(e1490)* V; *hmnIs102* [*col-56*pro:*tra-2*; *unc-122*pro:GFP]; *hmnEx2422* [*pkd-2*pro:GFP; *dat-1*pro:mApple; *rol-6(su1006)*] | 5C-5D |
| CHB4369 | *hmnIs82* [*grl-18*pro:GFP] II; *him-5(e1490)* V; *hmnIs102* [*col-56*pro:*tra-2*; *unc-122*pro:GFP] | 5F, Table S5C |
| CHB4375 | *hmnIs82* [*grl-18*pro:GFP] II; *him-5(e1490)* V; *hmnIs105* [*col-56*pro:*fem-3*; *unc-122*pro:GFP] | 5I, Table S5D |
| CHB3515 | *hmnIs47 [grl-18*pro:mApple] I; *him-5(e1490)* V | Table 1, Table S1 |
| CHB3916 | *hmnIs47 [grl-18*pro:mApple] I; *mab-3(hmn289)* II; *him-5(e1490)* V | Table 1 |
| CHB3716 | *hmnIs47 [grl-18*pro:mApple] I; *mab-3(mu15)* II; *him-5(e1490)* V | Table 1 |
| CHB4640 | *hmnIs82* [*grl-18*pro:GFP] II; *lep-2(hmn305)* IV; *him-5(e1490)* V | Table 1 |
| CHB3897 | *hmnIs82* [*grl-18*pro:GFP] II; *lep-2(ok900)* IV; *him-5(e1490)* V | Table 1 |
| CHB3887 | *hmnIs82* [*grl-18*pro:GFP] II; *him-5(e1490)* V; *lep-5(ny28)* X | Table 1 |
| CHB4186 | *hmnIs82* [*grl-18*pro:GFP] II; *him-5(e1490)* V; *jmjd-3.1(hmn304)* X | Table 1 |
| CHB4248 | *hmnIs82* [*grl-18*pro:GFP] II; *him-5(e1490)* V; *jmjd-3.1(gk384)* X | Table 1 |
| CHB4108 | *hmnIs82* [*grl-18*pro:GFP] II; *nfya-1(hmn316)* X | Table 1 |
| CHB4109 | *hmnIs82* [*grl-18*pro:GFP] II; *nfya-1(hmn317)* X | Table 1 |
| CHB4111 | *hmnIs82* [*grl-18*pro:GFP] II; *nfya-1(hmn319)* X | Table 1 |
| CHB4635 | *nfyb-1(tm4257)* II; *him-5(e1490)* V; *hmnIs78* [*grl-18*pro:GFP] X | Table 1 |
| CHB4606 | *nfyc-1(tm4264)* II; *him-5(e1490)* V; *hmnIs78* [*grl-18*pro:GFP] X | Table 1 |
| CHB4289 | *hmnIs82* [*grl-18*pro:GFP] II; *bed-3(hmn318)* IV; *him-5(e1490)* V | Table 1 |
| CHB4249 | *hmnIs82* [*grl-18*pro:GFP] II; *bed-3(sy705)* IV; *him-5(e1490)* V | Table 1 |
| CHB4400 | *hmnIs82* [*grl-18*pro:GFP] II; *bed-3(gk996)* IV; *him-5(e1490)* V | Table 1 |
| CHB4470 | *grl-18*(*hmn340*[*grl-18*-SL2-YFP:H2B + loxP]) *him-5(e1490)* V | S1G-S1J |
| CHB3992 | *hmnEx2223* [*col-56*pro:GFP; *dat-1*pro:mApple; *rol-6(su1006)*] | S2A |
| CHB4284 | *hmnIs82* [*grl-18*pro:GFP] II; *him-5(e1490)* V; *myEx696* [*pdf-1*pro:RFP; *unc-122*pro:GFP] | S3B |
| CHB4246 | *hmnIs82* [*grl-18*pro:GFP] II; *him-5(e1490)* V; *nfya-1(ok1174)* X; *myEx696* [*pdf-1*pro:RFP; *unc-122*pro:GFP] | S3B |
| CHB4360 | *him-5(e1490)* V; *kyIs53* [*odr-10*:GFP] X | S3D |
| CHB4352 | *him-5(e1490)* V; *kyIs53* [*odr-10*:GFP] *nfya-1(ok1174)* X | S3D |
| CHB4567 | *grl-18(hmn341) him-5(e1490)* V; *hmnEx2422* [*pkd-2*pro:GFP; *dat-1*pro:mApple; *rol-6(su1006)*] | S5B |
| CHB4583 | *grl-18(ok2845) him-5(e1490)* V; *hmnEx2422* [*pkd-2*pro:GFP; *dat-1*pro:mApple; *rol-6(su1006)*] | S5B |
| CHB3910 | *hmnIs47 [grl-18*pro:mApple] I; *him-8(e1489)* IV; *mab-23(gk664)* V | Table S1 |
| CHB3828 | *hmnIs47 [grl-18*pro:mApple] I; *him-8(e1489)* IV; *dmd-3(ok1327)* V | Table S1 |
| CHB3844 | *hmnIs47 [grl-18*pro:mApple] I; *dmd-5(ok1394)* II; *him-8(e1489)* IV | Table S1 |
| CHB3898 | *hmnIs47 [grl-18*pro:mApple] I; *dmd-6(gk287)* IV; *him-5(e1490)* V | Table S1 |
| CHB3768 | *hmnIs47 [grl-18*pro:mApple] I; *him-8(e1489)* IV; | Table S1 |

| | | |
|---|---|---|
| | *dmd-7(ok2276)* V | |
| CHB3769 | *hmnIs47 [grl-18*pro:mApple] I; *dmd-8(ok1294)* V | Table S1 |
| CHB3717 | *hmnIs47 [grl-18*pro:mApple] I; *dmd-9(ok1438)* IV; *him-5(e1490)* V | Table S1 |
| CHB3901 | *hmnIs47 [grl-18*pro:mApple] I; *him-8(e1489)* IV; *dmd-10(gk1125)* V | Table S1 |

### SUPPLEMENTAL TABLE S3

### Supp. Table S3. Plasmids generated in this study

| Plasmid | Description |
|---|---|
| pTT03.3 | *mir-228*pro:*fem-3* |
| pWF1 | *col-56*pro:GFP |
| pWF4 | *mir-228*pro:*tra-2* |
| pWF8 | *pkd-2*pro:GFP |
| pWF15 | *grl-18*pro:*grl-18*-gDNA:sfGFP |
| pWF16 | *grl-18*pro:sfGFP:*grl-18*-gDNA |
| pWF17 | *dat-1*pro:mApple |
| pWF22 | *col-56*pro:*fem-3* |
| pWF23 | *col-56*pro:*tra-2* |
| pWF53 | *col-56*pro:*nfya-1*-cDNA |
| pWF54 | *nfya-1*pro:*nfya-1*-cDNA |
| pWF107 | SapTrap_*grl-18*-SL2-YFP:H2B |
| pWF114 | *grl-18*pro:sfGFP:*grl-18*-gDNA(31-351) |
| pWF115 | *grl-18*pro:sfGFP:*grl-18*-gDNA(352-445) |
| pWF125 | pU6:*grl-18*-sgRNA #1 |
| pWF126 | pU6:*grl-18*-sgRNA #2 |

### SUPPLEMENTAL TABLE S4

### Table S4. Alleles generated in this study.

Upper case, exons; lower case; introns.

| Allele | Sequence | Notes |
|---|---|---|
| *hmn289* | GATGGGAAGCGTGTC[C>T]GTGACCCACACTGTG | R90C mutation |
| *hmn304* | CGGGAACGTGCGTCT[G>A]GTTTGCTGTGCCGTA | W831STOP mutation |
| *hmn305* | AAACGTGCGGAATTT[G>A]TATGGAGAACATTTT | C216Y mutation |
| *hmn316* | CCGCCAAACTGCAAG[g>a]ttagggagttttaca | splice site mutation |
| *hmn317* | CAAGCTCCGGCAAAG[C>T]GACCAATTCCGCCTA | R21STOP mutation |
| *hmn318* | AACAAGCTGCCGAG[T>G]GTCGAATTTGCCGGA | C144G mutation |
| *hmn319* | CCAGCTCCGGTTTCT[C>T]AATCACAACCACAGA | Q146STOP mutation |
| *hmn340* | TTCTTTCAACCTTGA[->atggctgtctcatcctactttcacctagttaactgcttgtcttaaaatctatgcttctctttagtatctaaaattttcctagaagcttacaagtatataaatggtctcttctcaataaaggttgtatatttattcatcttattgaatctgccatttcctcgtttttgcgagtttatataccttccaattttctttctattgtattttcaacttctaattttaattcagggaaactgctaccggtATGAGTAAAGGAGAAGAACTTTTCACTGGAGTTGTCCCAATTCTTGTTGAATTAGATGGTGATGTTAATGGGCACAAATTTTCTGTCAGTGGAGAGGGTGAAGGTGATGCAACATACGGAAAACTTACCCTTAAATTTATTTGCACTACTGGAAAACTACCTGTTCCATGGgtaagtttaaacatatatatactaactaaccctgattatttaaattttcagCCAACACTTGTCACTACTTTCGGTTATGGTCTTCAATGCTTCGCGAGATACCCAGATCATATGAAACGGCATGACTTTTTCAAGAGTGCCATGCCCGAAGGTTATGTACAGGAAAGAACTATATTTTTCAAAGATGACGGGAACTACAAGACACGTaagtttaaacagttcggtactaactaaccatacatatttaaattttcaggtGCTGAAGTCAAGTTTGAAGGTGATACCCTTGTTAATAGAATCGAGTTAAAAGGTATTGATTTTAAAGAAGATGGAAACATTCTCGGACACAAATTGGAATACAACTATAACTCACACAATGTATACATCATGGCAGACAAACAAAAGAATGGAATCAAAGTTgtaagtttaaacatgattttactaactaactaatctgatttaaattttcagAACTTCAAAATTAGACACAACATTGAAGATGGAAGCGTTCAACTAGCAGACCATTATCAACAAAATACTCCAATTGGCGATGGCCCTGTCCTTTTACCAGACAACCATTACCTGTCCTATCAATCTGCCCTTTCGAAAGATCCCAACGAAAAGAGAGACCACATGGTCCTTCTTGAGTTTGTAACAGCTGCtGGGATTACACATGGCATGGATGAACTATACGAATTCCCACCAAAGCCATCTGCCAAGGGAGCCAAGAAGGCCGCCAAGACCGTTACGAAGCCAAAGGACGGAAAGAAGAGACGTCATGCCCGTAAGGAATCATACTCCGTCTACATCTACCGTGTCCTCAAGCAAGTTCATCCAGACACTGGAGTTTCCTCCAAAGCCATGTCTATCATGAACTCTTTTGTCAACGATGTCTTCGAGCGTATTGCTGCTGAAGCATCCCGTCTTGCTCACTACAACAAGCGTTCCACAATCTCATCCCGCGAAATTCAGACCGCTGTCCGTCTGATCCTTCCAGGAGAGCTTGCCAAGCACGCCGTGTCTGAGGGAACCAAGGCCGTTACCAAGTACACTTCCAGCAAGTAGTGAgcggccgcaaggtaagtttaaaataacttcgtatagcatacattatacgaagttattttcagggtggcagcggaggtaccggcggtagtggaggcacg]ttcaaactttgatac | insertion of SL2-YFP:H2B + loxP + linker sequence immediately after the stop codon of the endogenous *grl-18* locus |
| *hmn341* | GAAAAGATGAAACATC[AACTCCAAAGTTCCCATTTGACCGAGATGAAAACGTAAACAAATGGACGAGTAAACGGAAAATTCGAGAAAAAGAAGAAATTAATTCAAAATGTAATAATCCGATTCTAAAGGATCTTATGGAAATGgtaatgagtcactggccaacagattccaagttaatattttttttagAAAATGACAACGTCTCCTTCAATATCGAAACAAATGATTTATTCAGCAGCAACCGAAATGTGGATGGGAAGGAATGTGAATGTGATATGTTCGAAACATTCATTTTCATATGTCGTTGTGACTTCTCCAATTTTTTGTGAACACAGGAAAAAGGCGTTGACATGCTTCGTTTTCTTTCAACCTTGAttcaaactttgatactctttttttattgttagggtgctttttgataataaattattatttattccatgttatactaaaatgtatgactgaaaatctactttttctgcttgaactcagcggatatgaaaccttttcaaattttgttgtaaaattctgaaaaaacacaaaatttgcaacaaaatggctacggggatttttgaaaaagtttgtttgaacccagatattagttttaaaaaattggaaaatctactaattttgcatataaatggtcggttttagtcaatctcatctcaagctgtgcactgtgctagaaatgtgcaaaaaaaaagcaaaaattagacgaaaattgaaaaatgttagacaaaaaagttttttaattcagaaaaaattttaactattttaatttttctttctgttctgccaatatgtttcctctagagtcctgctaatcgttttgatttttttgaaaatttctattaaaaatttcacagttattttagtcatttagggctaaaaacccacatacgctacttcaaagtattcgattcataattctatttttatttatttgtcaaaactacacaagcagctttcaattagtc>-]gaaaattctagtcatc | deletion spanning Grl domain of the endogenous *grl-18* locus |

| | | |
|---|---|---|
| *syb6299* | ATATTTTTCCCTCAA[CTT>TTG]GGATTCG[gtaaactttaattgtaactccctggaaagtccatgaattaaaattcagGT>GATCCATGAGCAAAGGAGAAGAACTTTTCACTGGAGTTGTCCCAATTCTTGTTGAATTAGATGGTGATGTTAATGGGCACAAATTTTCTGTCCGTGGAGAGGGTGAAGGTGATGCTACAAACGGAAAACTCACCCTTAAATTTATTTGCACTACTGGAAAACTACCTGTTCCGTGGCCAACACTTGTCACTACTCTGACCTATGGTGTTCAATGCTTTTCCCGTTATCCGGATCACATGAAACGGCATGACTTTTTCAAGAGTGCCATGCCCGAAGGTTATGTACAGGAACGCACTATATCTTTCAAAGATGACGGGACCTACAAGACGCGTGCTGAAGTCAAGTTTGAAGGTGATACCCTTGTTAATCGTATCGAGTTAAAGGGTATTGATTTTAAAGAAGATGGAAACATTCTTGGACACAAACTCGAGTACAACTTTAACTCACACAATGTATACATCACGGCAGACAAACAAAAGAATGGAATCAAAGCTAACTTCAAAATTCGCCACAACGTTGAAGATGGTTCCGTTCAACTAGCAGACCATTATCAACAAAATACTCCAATTGGCGATGGCCCTGTCCTTTTACCAGACAACCATTACCTGTCGACACAATCTGTCCTTTCGAAAGATCCCAACGAAAAGCGTGACCACATGGTCCTTCTTGAGTTTGTAACTGCTGCTGGGATTACACATGGCATGGATGAGCTCTACAAAGGA]TCCATGAACTGCCAATGCCAAAATTCGTGT[AG>TC]CTCTCCACCGGCACAA | removal of the first intron, insertion of sfGFP, and synonymous mutations to the endogenous *grl-18* locus |
